# RegFormer: A Single-Cell Foundation Model Powered by Gene Regulatory Hierarchies

**DOI:** 10.1101/2025.01.24.634217

**Authors:** Luni Hu, Hua Qin, Yilin Zhang, Yi Lu, Ping Qiu, Zhihan Guo, Lei Cao, Wenjian Jiang, Qianqian Chen, Yanbang Shang, Tianyi Xia, Ziqing Deng, Xun Xu, Shuangsang Fang, Yuxiang Li, Yong Zhang

## Abstract

Single-cell RNA sequencing (scRNA-seq) enables high-resolution profiling of cellular diversity, but current computational models often fail to incorporate regulatory priors, handle data sparsity, or efficiently process long gene sequences. Here, we present RegFormer, a foundation model that integrates gene regulatory networks (GRNs) with Mamba-based state-space modeling, overcoming the scalability and context-length limitations of Transformer architectures. RegFormer encodes each gene through dual embeddings, a value embedding for quantitative expression and a token embedding for regulatory identity, organized within a GRN-guided gene order to capture both expression dynamics and hierarchical regulation. Pretrained on 26 million human single cells spanning 45 tissues and diverse biological contexts, RegFormer achieves superior scalability and biological fidelity. Across comprehensive benchmarks, it consistently outperforms state-of-the-art single-cell foundation models (scGPT, Geneformer, scFoundation, and scBERT), delivering higher clustering accuracy, improved batch integration, and more precise cell type annotation. RegFormer also reconstructs biologically coherent GRNs, accurately models transcriptional responses to genetic perturbations, and enhances drug response prediction across cancer cell lines. By combining regulatory priors with efficient long-sequence Mamba modeling, RegFormer establishes a biologically grounded and scalable framework for single-cell representation learning, enabling deeper mechanistic insight into gene regulation and cellular state transitions.

## Introduction

Single-cell RNA sequencing (scRNA-seq) has revolutionized our understanding of cellular heterogeneity, gene expression dynamics, and the complex regulatory networks that underpin cellular processes^1–3^. By enabling the detailed characterization of individual cells, scRNA-seq has unveiled key biological mechanisms involved in tissue development^4^, disease progression^5^, and responses to environmental stimuli^6^. Large-scale initiatives like the Human Cell Atlas exemplify the growing scope of single-cell transcriptomics, providing comprehensive maps of cellular diversity across tissues and disease contexts^7^. Despite these transformative insights, computational methods for analyzing scRNA-seq data remain fragmented, with models often tailored for specific tasks and lacking scalability across diverse datasets and cellular states^8–10^, underscoring the need for a unified, integrated analytical framework capable of performing multiple tasks simultaneously with higher accuracy and consistency.

To address this gap, recent advances have led to the development of single-cell foundation models, such as scGPT^11^, Geneformer^12^, scFoundation^13^, and scBERT^14^, which aim to provide a general-purpose solution for the analysis of scRNA-seq data. These models leverage deep learning techniques, which have proven effective at capturing complex patterns within high-dimensional, sparse scRNA-seq datasets. By learning representations of gene expression profiles, these models can achieve strong performance in tasks like cell annotation and genetic perturbation prediction. However, despite their success, these models typically rely on natural language processing (NLP) algorithms^15, 16^, which are inherently designed for sequential data, such as text or genomic sequences. While these approaches are beneficial in certain contexts, they do not fully address the unique challenges posed by scRNA-seq data, specifically the unordered nature of gene expression measurements across individual cells. Unlike traditional sequence data, where elements (such as nucleotides or words) follow a fixed order, scRNA-seq data lacks this inherent structure, which makes it difficult for sequence-based models to capture the complex, non-linear interactions between genes and the hierarchical relationships that define cellular states.

To overcome these limitations, biological prior knowledge, particularly that derived from gene regulatory networks (GRNs), has been increasingly incorporated into scRNA-seq analysis^17–19^. GRNs represent the intricate regulatory relationships between genes and provide a framework for understanding how gene expression is regulated within cellular contexts. These networks capture key biological insights that are not immediately apparent from raw gene expression data alone, such as transcriptional hierarchical dependencies, and spatial-temporal regulation of gene activity^20, 21^. GRN-based priors enable models to capture regulatory mechanisms more faithfully and enhance biological relevance. This not only enhances the accuracy of predictions but also provides richer, more biologically grounded insights into the underlying regulatory mechanisms that govern cellular behavior and function. The integration of GRNs allows for the modeling of hierarchical dependencies between genes, which is critical for understanding cellular dynamics, differentiation, and disease mechanisms.

Mamba Blocks provide an innovative solution for analyzing high-dimensional, sparse datasets by enabling flexible representation learning and hierarchical modeling of gene interactions. Unlike traditional transformer architectures, which often struggle with computational complexity, Mamba Blocks are specifically engineered to effectively address the intricacies of scRNA-seq data^22^.Existing models such as DGRNA^23^, Orthrus^24^, and SC-MAMBA2^25^, which are also based on Mamba architecture, primarily focus on scRNA-seq data and broader RNA biology tasks. DGRNA excels at modeling RNA sequences, while Orthrus specializes in splicing dynamics. However, these models may not fully capture the complexities of scRNA-seq data, which are essential for understanding cellular states. Although SC-MAMBA2 performs well in single-cell analysis, it tends to emphasize gene-gene interactions without integrating biological prior knowledge.By leveraging hierarchical insights from gene regulatory networks, Mamba Blocks enhance our understanding of cellular behavior and function, effectively addressing the need for more biologically grounded computational models in single-cell research.

Here, we present RegFormer, a foundation model tailored for scRNA-seq analysis. RegFormer integrates GRNs with a Mamba-based state-space architecture optimized for high-dimensional and sparse expression data. This design enables efficient modeling of long-range dependencies across gene sequences, capturing both local interactions and global regulatory hierarchies. Through generative pretraining on 26 million human cells, RegFormer learns gene expression dynamics together with the hierarchical structure of gene regulation, enhancing biological fidelity. By incorporating GRN-guided priors, it effectively models transcriptional dependencies within regulatory hierarchies, providing mechanistic insights into gene control and cellular behavior. Across diverse benchmarks, including cell annotation, GRN reconstruction, genetic perturbation prediction, and drug response modeling, RegFormer consistently outperforms existing foundation models such as scGPT and Geneformer. Together, these advances position RegFormer as a scalable framework for decoding complex transcriptional programs and advancing single-cell biology.

## Results

### RegFormer leverages regulatory priors for single-cell modeling with Mamba

RegFormer is a foundation model that integrates regulatory priors into single-cell transcriptome modeling through the combination of gene regulatory networks (GRNs) and Mamba sequence modeling (**Fig. 1A**). For large-scale pretraining, we curated a harmonized collection of 26 million human single cells spanning 45 tissues, multiple sequencing technologies, and diverse developmental and disease contexts (**Supplementary Fig. S1B-E**). Data preprocessing followed a unified pipeline: raw matrices were imported as AnnData objects, filtered for low-quality genes and cells, normalized, log-transformed, and written into LMDB for efficient large-scale training **(Supplementary Fig. S1A**).

**Figure 1.**
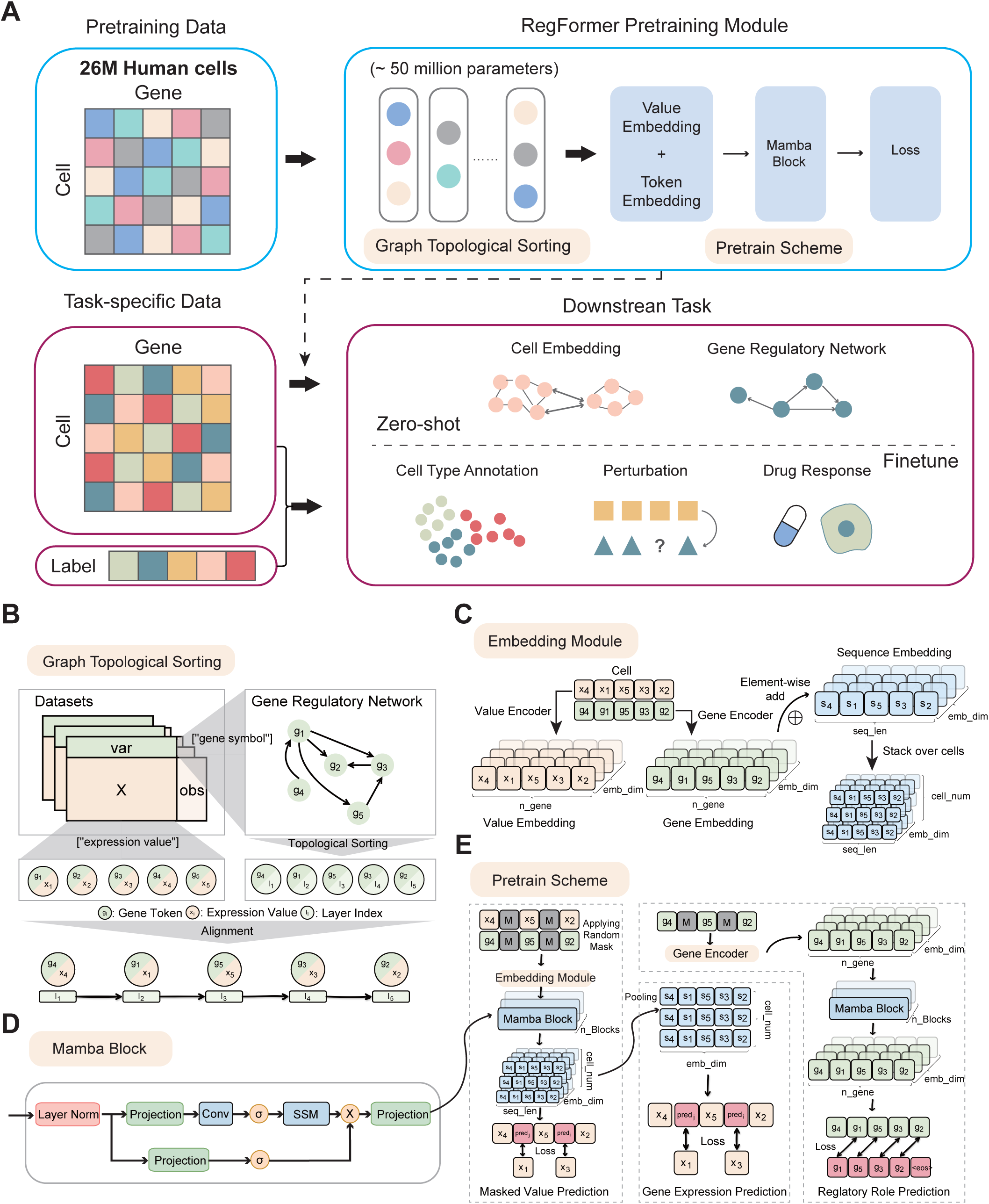
Overview of the RegFormer framework. **(A)** The RegFormer pretraining and finetuning workflow. Large-scale pretraining on 26 million human cells was performed using graph topological sorting, value/token embeddings, and Mamba blocks, followed by downstream tasks including cell embedding, gene regulatory network (GRN) inference, cell type annotation, perturbation modeling, and drug response prediction. **(B)** Graph topological sorting strategy for constructing regulatory dependencies from gene expression matrices. **(C)** The embedding module design integrating value embeddings and gene encoders into sequence embeddings. **(D)** Architecture of the Mamba block within the encoder. **(E)** Pretraining schemes including masked value prediction, gene expression prediction, and regulatory role prediction.

To incorporate biological prior knowledge, we constructed GRNs by linking transcription-factor (TF) motifs to their downstream targets using curated motif-to-gene mappings. The networks were refined via Node2Vec embeddings and iterative edge removal until a directed acyclic graph (DAG) was obtained (**Supplementary Fig. S2A**). This process retained > 92 % of edges while preserving degree and path-length distributions (**Supplementary Fig. S2B-F**). The final network contained 13.4 % TF nodes and 86.6 % target genes arranged across hierarchical regulatory depths (**Supplementary Fig. S3A-C**). Genes were then topologically sorted along this hierarchy to align expression matrices with regulatory order (**Fig. 1B**)..

Within this GRN-ordered framework, RegFormer encodes each gene using dual embeddings: a value embedding representing quantitative expression and a token embedding representing the gene’s regulatory identity (**Fig. 1C**). These embeddings are processed by Mamba blocks, which combine multi-layer perceptrons, convolutional operations, and state-space modules to capture both local interactions and long-range dependencies (**Fig. 1D**). During pretraining, RegFormer is optimized through three complementary self-supervised objectives, masked-value prediction, gene expression prediction, and regulatory role prediction, which together enable the model to learn quantitative variation, expression reconstruction, and hierarchical regulatory relationships (**Fig. 1E**). Together, these components form a unified framework that integrates biological priors with sequence modeling to learn the regulatory principles underlying single-cell gene expression.

### Mechanistic validation of GRN guidance and architectural design

We further evaluated how regulatory priors and architectural design influence RegFormer’s learning behavior through systematic ablation and functional analyses. All models were trained on **1 million blood cells** and **evaluated across eight validation datasets** to ensure robust and comparable assessment. (**Supplementary Figs. S4-S13**). RegFormer embeddings accurately captured transcriptional hierarchy and maintained proximity between transcription factors and their targets. Compared with randomly initialized or MLM-based controls, RegFormer achieved higher transcription factor classification accuracy and stronger correlations between gene embeddings and regulatory degree, indicating that the learned representations closely align with the underlying GRN topology (**Supplementary Figs. S5-S6**). Functionally, reconstructed GRNs derived from RegFormer embeddings exhibited greater biological coherence: across eight blood validation datasets, GRN-guided models yielded more enriched pathways and higher cross-dataset functional similarity than non-GRN variants (**Supplementary Fig. S7**), demonstrating that regulatory priors improve biological fidelity.

Ablation experiments further elucidated the contribution of each architectural component. Replacing Mamba blocks with Transformer layers weakened cell-level representation and reduced preservation of gene-level regulatory relationships (**Supplementary Fig. S8**), indicating that the state-space formulation more effectively captures hierarchical transcriptomic dependencies. Across blood validation datasets, GRN-guided RegFormer consistently achieved higher Average Silhouette Width (ASW) scores than the Random variant, confirming the advantage of incorporating regulatory priors (**Supplementary Fig. S9A**). Removing the token embedding and retaining only value embeddings led to lower clustering performance, suggesting that regulatory context encoded by gene tokens is essential for representing cell-type structure (**Supplementary Fig. S9B**). Substituting the regression-based value prediction with binary classification (BinCls) also decreased ASW scores, indicating that modeling gene expression as a continuous variable captures quantitative variation more accurately (**Supplementary Fig. S9C**). When individual pretraining objectives, including masked-value prediction (MVP), gene-expression prediction (GEPC), or regulatory-role prediction (TOPO), were removed, model performance consistently declined across datasets, with the largest decrease observed when the TOPO objective was omitted (**Supplementary Fig. S9D**).

Similarly, shuffling or removing GRN edges substantially reduced ASW performance and weakened degree correlations and reconstruction precision (**Supplementary Fig. S10**), underscoring the essential contribution of authentic regulatory topology to model stability and biological coherence. Notably, extending the input sequence from 1.2k to 10k genes further improved clustering resolution and functional consistency (**Supplementary Fig. S11**). The longer input sequence provided a more comprehensive regulatory context and enabled the model to capture long-range dependencies across transcriptional modules, thereby enhancing both biological insight and scalability. In addition, compared with graph neural network frameworks, RegFormer achieved higher accuracy at both the gene and cell levels (**Supplementary Fig. S12**).

Together, GRN-guided gene ordering, dual-embedding design, and Mamba-based long-sequence modeling act synergistically to allow RegFormer to capture cross-hierarchical regulatory relationships within an expanded gene space, establishing a biologically grounded and scalable framework for single-cell representation learning.

### RegFormer achieves superior single-cell embeddings across tissues

Single-cell foundation models have emerged as powerful tools for representing, classifying, and analyzing the complex information embedded within single-cell datasets. Using the BioLLM benchmarking framework^26^, we systematically evaluated the representation capacity of RegFormer against state-of-the-art single-cell foundation models, including scGPT, GeneFormer, scFoundation, and scBERT, across multiple human tissue datasets (**Fig. 2**). RegFormer consistently achieved higher Average Silhouette Width (ASW) scores, indicating superior separation of biologically distinct cell populations. In the human lung dataset, representing a discretized single-cell transcriptome, RegFormer accurately resolved epithelial, endothelial, and immune lineages, effectively capturing fine-grained intra-tissue heterogeneity (**Fig. 2A**). In the bone marrow dataset, representing a continuous single-cell trajectory, RegFormer outperformed other models in distinguishing early myeloid and lymphoid progenitors (**Fig. 2B**). In the dendritic cell dataset, RegFormer maintained subtype consistency across batches while achieving high cell-type ASW and low batch ASW values, demonstrating strong batch invariance (**Fig. 2C**). In the pancreas dataset, the model robustly separated endocrine, exocrine, and stromal compartments, even in the presence of substantial technical variation (**Fig. 2D**).

**Figure 2.**
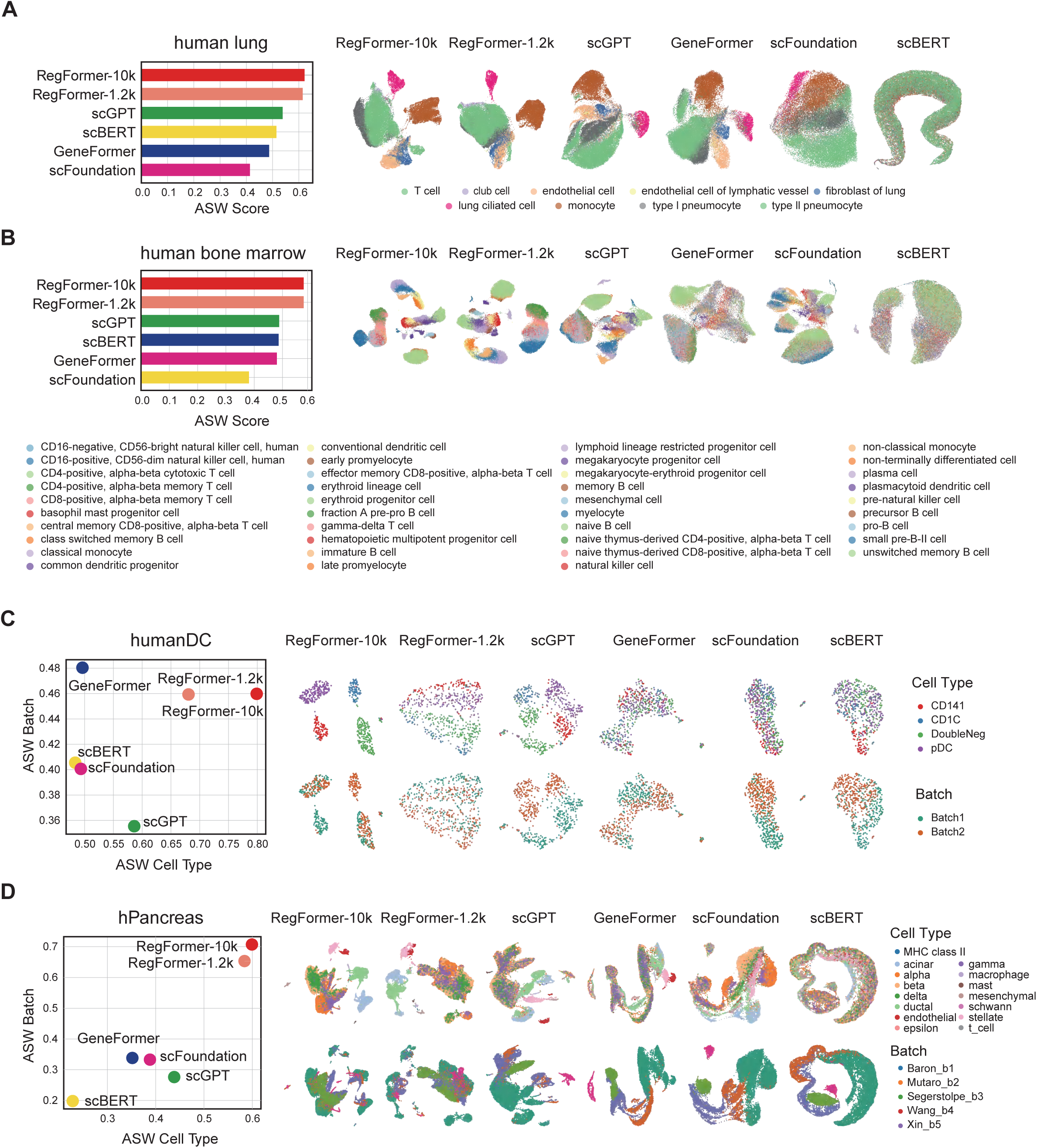
Benchmarking RegFormer Embedding against single-cell foundation models. **(A)** ASW scores and UM AP projection of cell types in the human lung dataset. **(B)** ASW scores and UMAP projection of cell types in the human bone marrow dataset. **(C)** ASW scores and UMAP projection of cell type and batch information in the human dendritic cell (humanDC) dataset. **(D)** ASW scores and UMAP projection of cell type and batch information in the human pancreas (hPancreas) dataset.

Performance analyses across varying numbers of highly variable genes (HVGs) further confirmed the scalability and stability of RegFormer embeddings (**Supplementary Fig. S13**). Both RegFormer-1.2k and RegFormer-10k maintained consistently high ASW scores across HVG subsets, whereas other models showed marked performance declines as the number of features decreased. The broader-context 10k variant provided additional gains, indicating that longer gene sequences supply a more complete regulatory context and enhance the model’s ability to capture gene regulatory information.

### RegFormer enhances cell type annotation performance across diverse datasets

To further evaluate the downstream application of RegFormer, we benchmarked its performance on cell type annotation tasks across six representative datasets, in comparison with leading single-cell foundation models. RegFormer-10k and RegFormer-1.2k consistently achieved higher Macro-averaged F1 score (Macro-F1) and comparable accuracy values than all baseline models, demonstrating more precise and generalizable cell type classification (**Fig. 3A**). In the Zheng68k dataset, RegFormer accurately reconstructed the fine-grained immune landscape, effectively distinguishing closely related CD4 and CD8 T-cell subsets (**Fig. 3B-C**). Across additional tissues including blood, liver, small intestine, kidney, and M.S., RegFormer generated the most coherent UMAP embeddings and exhibited the highest diagonal concentration in confusion matrices, indicating faithful label recovery and minimal cross-type confusion (**Supplementary Fig. S14**).

**Figure 3.**
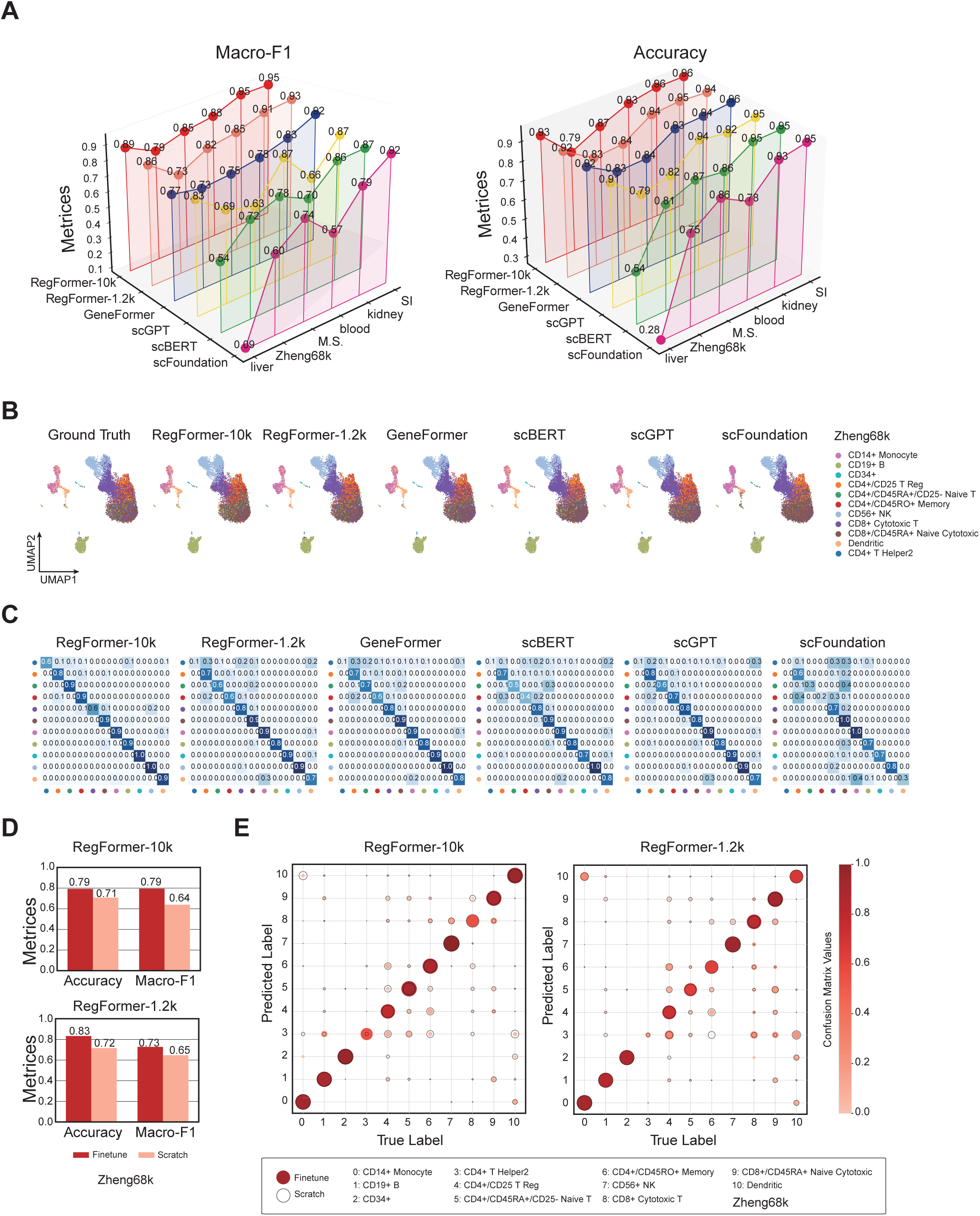
Cell type annotation performance of RegFormer. **(A)** Macro-F1 and accuracy across seven benchmark datasets, shown for RegFormer-10k, RegFormer-1.2k, Geneformer, scBERT, scGPT, and scFoundation. **(B)** UMAP projections of cell type annotations on the Zheng68k dataset, comparing ground truth with predictions from different models. **(C)** Confusion matrices of predicted versus true labels on Zheng68k across different models. **(D)** Comparison of RegFormer-10k and RegFormer-1.2k in accuracy and Macro-F1 under scratch training and finetuning settings on Zheng68k dataset. **(E)** Dot plots showing correspondence between predicted and true cell type labels for RegFormer-10k (left) and RegFormer-1.2k (right), with dot size proportional to predicted label proportion, solid circles representing finetune, and open circles representing scratch training.

In quantitative analyses, RegFormer achieved the highest macro-average precision (AP) across all benchmark datasets (**Supplementary Fig. S15A**). The model maintained stable predictive performance across cell types of varying abundance and showed stronger recognition ability for rare cell populations (**Supplementary Fig. S15B**). When finetuned on downstream cell type annotation tasks, both RegFormer-10k and RegFormer-1.2k achieved higher Macro-F1 and accuracy compared with scratch training (**Fig. 3D and Supplementary Fig. S16A**). Confusion matrix comparisons further showed improved subtype classification consistency after finetuning (**Supplementary Fig. S16B**), and cell type–wise precision analyses demonstrated consistent performance gains across different cell categories (**Supplementary Fig. S16C**).

Together, these results demonstrate that RegFormer provides accurate, scalable, and generalizable cell type annotation across heterogeneous single-cell datasets, with the extended 10k variant further improving fine-grained resolution and rare cell type recognition through broader gene coverage and regulatory context integration.

### RegFormer reconstructs biologically coherent gene regulatory networks

We next evaluated the ability of RegFormer to reconstruct gene regulatory networks (GRNs) from single-cell gene expression data. Gene embeddings derived from the pretrained model were used to compute pairwise cosine similarity between genes, and the resulting similarity matrices were interpreted as putative regulatory connections between transcription factors and their targets (**Fig. 4A**). Applied to the human lung dataset (**Fig. 4B**), RegFormer-inferred GRNs exhibited higher functional similarity than those generated by existing single-cell foundation models (**Fig. 4C**). RegFormer also achieved the highest Gene Ontology (GO) enrichment counts across a range of clustering resolutions (**Fig. 4D**), indicating that its embeddings better preserve biologically relevant gene interactions.

**Figure 4.**
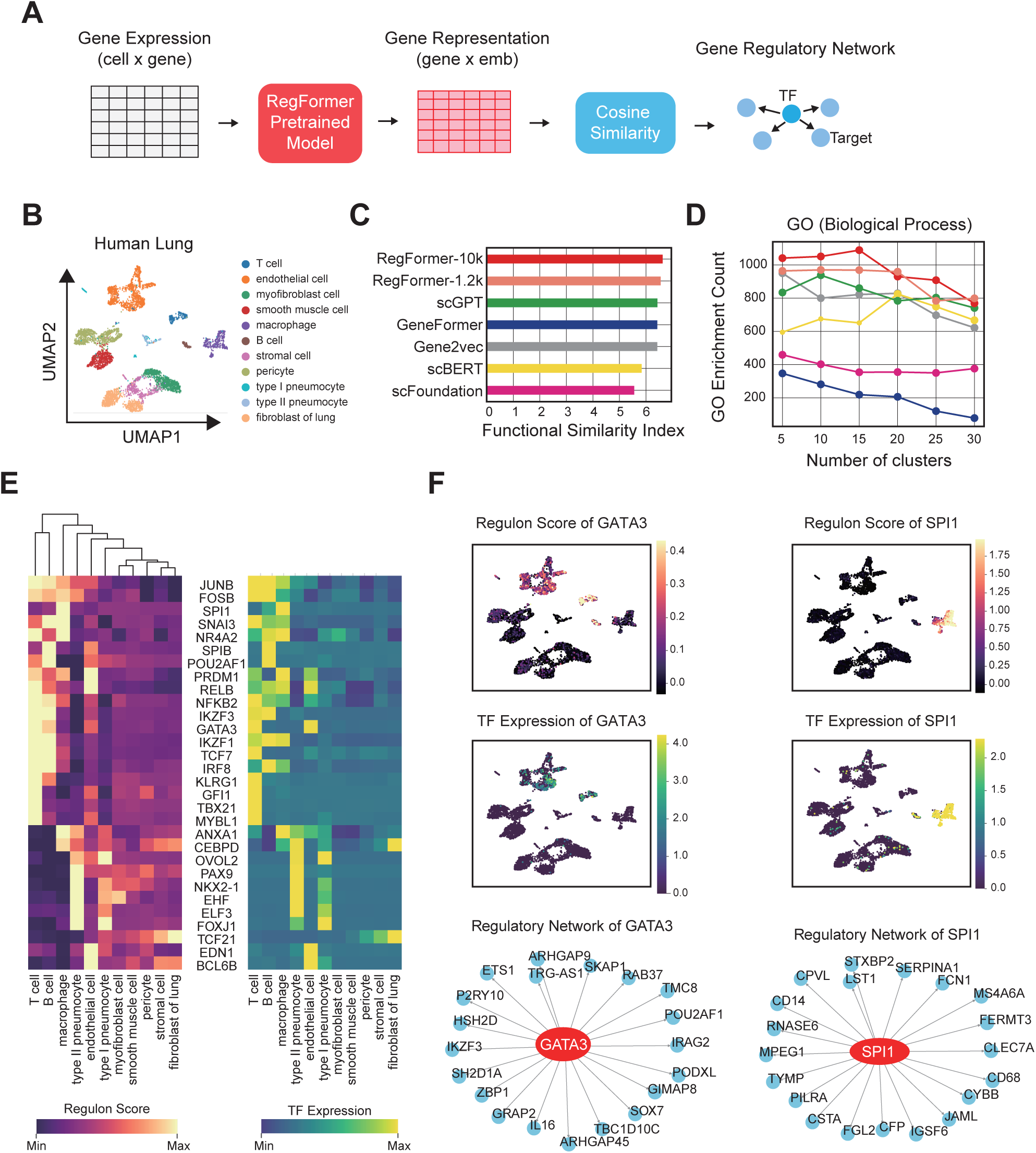
Gene regulatory network inference with RegFormer. **(A)** Workflow for GRN construction from gene expression profiles, using RegFormer embeddings and cosine similarity to infer regulatory links. **(B)** UMAP projection of cell types in the human lung dataset. **(C)** Functional similarity index of inferred GRNs comparing RegFormer with existing single-cell foundation models. **(D)** Gene Ontology (GO, Biological Process) enrichment analysis across different numbers of gene clusters, comparing results from RegFormer and other single-cell foundation models. **(E)** Heatmaps showing regulon activity scores and transcription factor (TF) expression patterns across diverse lung cell types. **(F)** Case studies of GATA3 and SPI1 regulons, including UMAP visualizations of regulon scores (top), TF expression (middle), and the corresponding inferred GRN subnetworks (bottom).

Analysis of regulon activity revealed a strong correspondence between transcription factor expression and inferred regulon scores across diverse lung cell types (**Fig. 4E**). Case studies of canonical regulators further supported the biological validity of the inferred networks (**Fig. 4F and Supplementary Fig. S17**). The regulon activity and expression of GATA3 were predominantly enriched in T cells, consistent with its established role as a master regulator of T-cell differentiation. In contrast, SPI1 (PU.1) activity and expression were concentrated in macrophage and monocyte-associated clusters, in line with its key role in myeloid lineage commitment. The reconstructed GATA3 and SPI1 regulons included canonical immune-related targets such as IL16, POU2AF1, CD14, and CLEC7A, accurately reflecting their roles in adaptive and innate immune regulation (**Fig. 4F**).

Comparisons between the input and reconstructed GRNs further demonstrated stronger cross-cell-type correlations among regulatory edges and improved concordance between transcription factors and their predicted targets (**Supplementary Fig. S18A-B**). Functional enrichment analyses revealed that the reconstructed networks provided enhanced representation of biologically relevant pathways, particularly those related to immune activation, morphogenesis, and neurogenesis (**Supplementary Fig. S18C-D**). Collectively, These findings indicate that RegFormer enables the reconstruction of biologically coherent and functionally meaningful GRNs, providing a robust framework for linking single-cell gene expression profiles to underlying regulatory mechanisms.

### RegFormer enables accurate modeling of genetic perturbations

Building on its ability to reconstruct static gene regulatory architectures, we next examined whether RegFormer embeddings could also capture dynamic transcriptional responses under genetic perturbations. By integrating pretrained gene embeddings with the GEARS framework, RegFormer predicts post-perturbation expression states from unperturbed profiles, modeling how transcriptional programs shift in response to targeted perturbations **(Fig. 5A)**.

**Figure 5.**
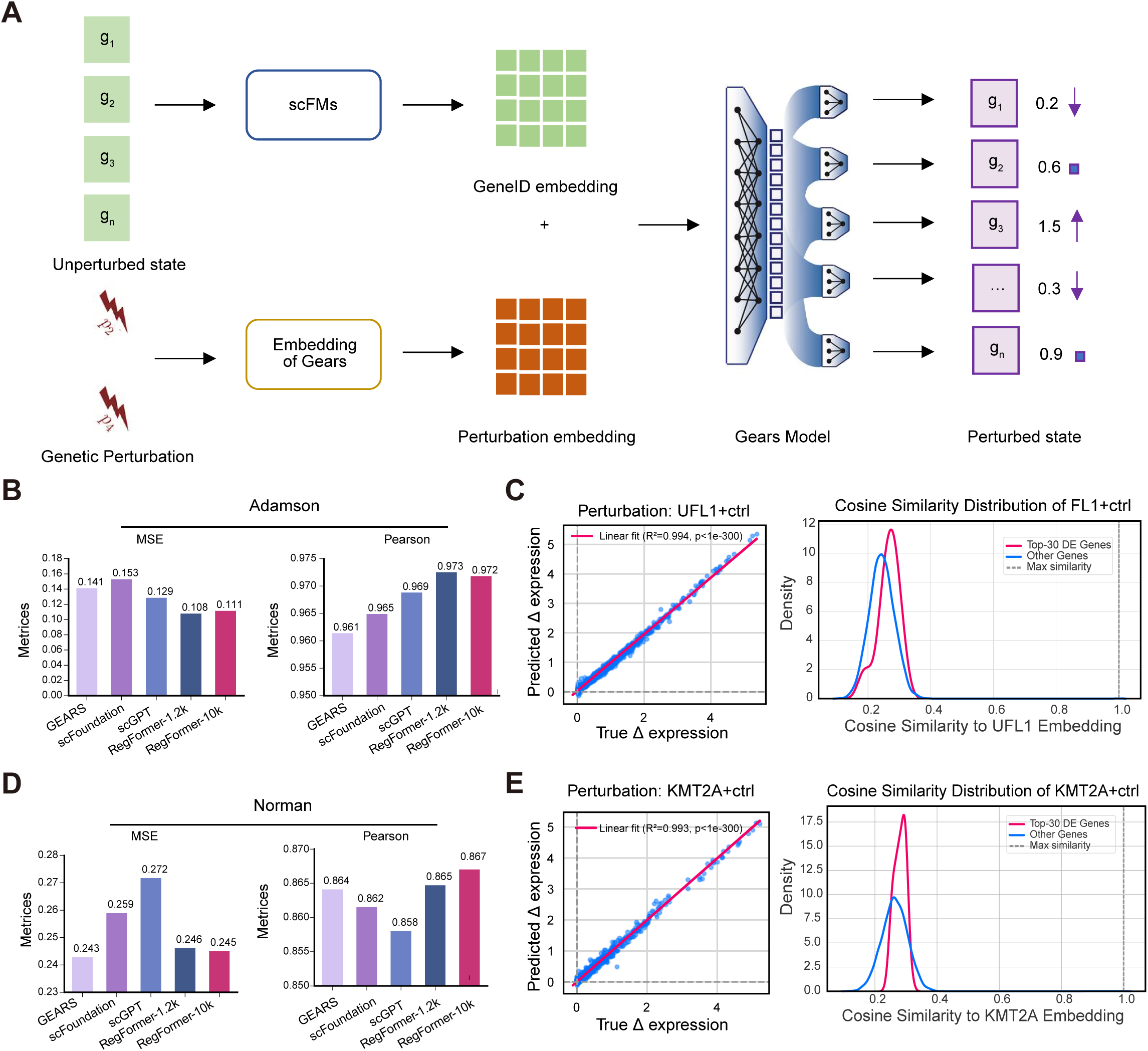
Perturbation modeling with RegFormer embeddings. **(A)** Schematic of perturbation prediction using embeddings from single-cell foundation models combined with the Gears model, integrating GeneID embeddings and perturbation embeddings to predict perturbed states. **(B)** Prediction performance on the Adamson CRISPR perturbation dataset, evaluated by mean squared error (MSE, left) and Pearson correlation (right), comparing GEARS, scFoundation, scGPT, RegFormer-1.2k, and RegFormer-10k. **(C)** Example perturbation UFL1+ctrl: scatter plot of predicted versus true Δ expression (left) and gene embedding cosine similarity between the perturbed TF and differentially expressed (DE) genes or other genes (right). **(D)** Prediction performance on the Norman CRISPR perturbation dataset, assessed by MSE (left) and Pearson correlation (right). **(E)** Example perturbation KMT2A+ctrl: scatter plot of predicted versus true Δ expression (left) and gene embedding cosine similarity between the perturbed TF and differentially expressed (DE) genes or other genes (right).

Across both the Adamson and Norman CRISPR perturbation datasets, RegFormer achieved the lowest mean squared error (MSE) and highest Pearson correlation among all single-cell foundation models tested, including scFoundation and scGPT (**Fig. 5B, D**). Predicted versus observed Δ-expression values showed near-perfect linear correspondence (R²>0.97) for representative perturbations such as UFL1 and KMT2A, confirming that RegFormer embeddings effectively encode regulatory dependencies (**Fig. 5C, E and Supplementary Fig. S19**). Cosine similarity analysis further revealed that differentially expressed genes exhibited higher embedding proximity to the perturbed transcription factor than nonresponsive genes, suggesting that RegFormer effectively captures the regulatory dependencies driving transcriptional responses (**Fig. 5C, E and Supplementary Fig. S19**).

Together, these findings demonstrate that RegFormer embeddings provide a unified representation that generalizes from static GRN topology to dynamic perturbation effects, enabling accurate modeling of transcriptional responses across diverse perturbation contexts.

### RegFormer improves drug response prediction across diverse cancer cell lines

Having demonstrated its ability to model both regulatory structure and perturbation dynamics, we next evaluated whether RegFormer embeddings could improve drug response prediction across cancer cell lines. We integrated RegFormer-derived gene expression embeddings with molecular graph representations of drug structures using a uniform graph convolutional network to predict IC50 values (**Fig. 6A**). Across multiple benchmarks, RegFormer achieved the highest Pearson Correlation Coefficient (PCC) and Spearman Rank Correlation Coefficient (SRCC) among all compared models, including DeepCDR, GeneFormer, scGPT, scBERT and scFoundation (**Fig. 6B-D and and Supplementary Fig. S20A-B**). Predicted versus observed IC50 values showed strong linear correspondence across both drugs and cancer types (**Fig. 6C**), and maintained high precision for the top five best-predicted compounds (**Fig. 6D**). In leave-drug-out blind tests, RegFormer consistently outperformed DeepCDR, achieving higher PCC across nearly all unseen drugs (**Fig. 6E and Supplementary Fig. S20C**).

**Figure 6.**
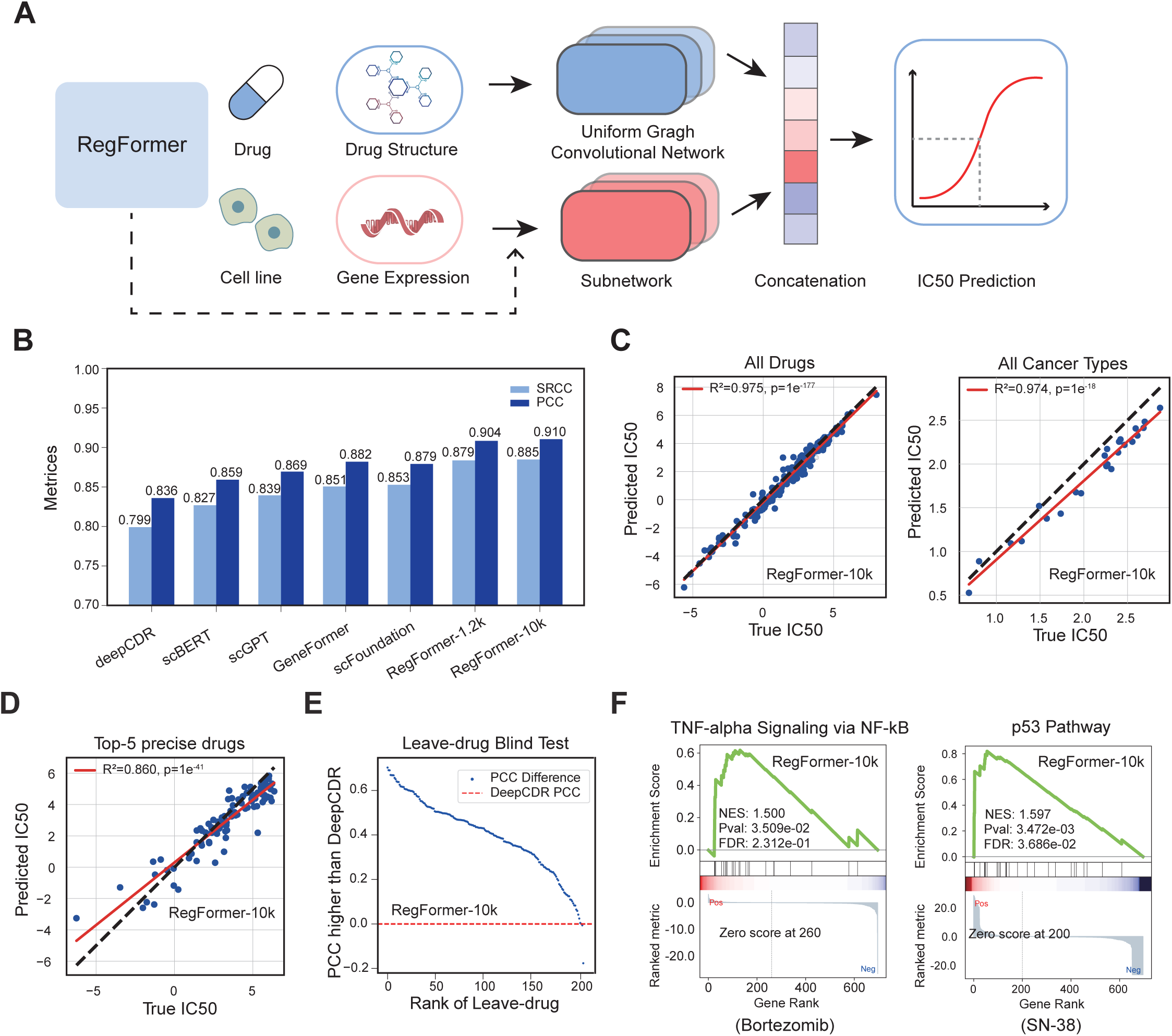
Drug response prediction using RegFormer. **(A)** Overview of the IC50 prediction framework. RegFormer embeddings of cell line gene expression and precomputed embeddings of drug structure are integrated through a uniform graph convolutional network to predict drug sensitivity (IC50). **(B)** Comparison of prediction performance across models, including deepCDR, scBERT, scGPT, Geneformer, scFoundation, RegFormer-1.2k, and RegFormer-10k, evaluated by Spearman (SRCC) and Pearson (PCC) correlation coefficients. **(C)** Predicted versus true IC50 values for all drugs (left) and across all cancer types (right), showing high correlation between predicted and observed responses. **(D)** Predicted versus true IC50 values for the top five most accurately predicted drugs, demonstrating strong linearity. **(E)** Leave-drug-out blind test, where RegFormer consistently achieves higher PCC than DeepCDR across unseen drugs. **(F)** Pathway-level interpretation of drug responses, highlighting TNF-α signaling via NF-κB (Bortezomib) and p53 pathway (SN-38), with normalized enrichment scores (NES) and FDR values shown.

Beyond numerical accuracy, pathway-level analyses revealed that RegFormer captured biologically meaningful mechanisms of drug sensitivity. For example, Bortezomib responses were associated with TNF-α signaling via NF-κB^27^, while SN-38 responses were linked to p53-mediated apoptosis^28^ (**Fig. 6F**). These results indicate that RegFormer embeddings significantly improve predictive performance in drug response prediction.

## Discussion

The introduction of RegFormer marks a major advance in single-cell RNA sequencing (scRNA-seq) analysis, addressing long-standing challenges in data sparsity, biological relevance, and scalability. By integrating gene regulatory networks (GRNs) within a Mamba-based architecture, RegFormer bridges biological knowledge with efficient long-sequence modeling, setting new benchmarks across key analytical tasks including cell annotation, GRN reconstruction, genetic perturbation prediction, and drug response modeling.

A central innovation of RegFormer is its use of GRNs to model hierarchical regulatory relationships fundamental to cellular identity. Unlike prior models such as scBERT and GeneFormer, which either lacked biological priors or were constrained by limited context windows, RegFormer orders genes according to GRN-guided topology, allowing the model to learn regulation in a biologically meaningful sequence. Performance remains sensitive to the quality and completeness of GRN priors, incomplete or noisy networks may distort ordering or weaken hierarchical signal propagation. Future work may employ context-specific or probabilistic GRNs to enhance adaptability across tissues. To ensure hierarchical consistency, cycles in GRNs are removed to form a directed acyclic graph (DAG), which preserves most structure but may omit feedback loops important for regulation. Developing differentiable or dynamic cycle-handling mechanisms could mitigate this limitation.

Architecturally, RegFormer represents a departure from Transformer-based models such as scGPT, GeneFormer, and scFoundation, which rely on quadratic attention mechanisms that limit both scalability and effective context length (**Supplementary Table S2**). In contrast, RegFormer’s Mamba Blocks adopt a state-space formulation with linear time complexity, enabling efficient modeling of extended gene sequences encompassing up to 10,000 genes. This formulation preserves long-range regulatory dependencies across transcriptional modules that are often attenuated or truncated in attention-based Transformers, thereby enhancing the model’s ability to represent coherent regulatory programs and global transcriptional coordination. Future work will extend benchmarking to full-transcriptome contexts, enabling comprehensive assessment of Mamba’s genome-scale modeling capacity and its potential to capture hierarchical regulatory organization across the entire transcriptome.

Empirical evaluations demonstrate that RegFormer achieves superior performance across diverse human tissues and datasets. In cell annotation tasks, it consistently outperforms existing models, yielding more precise classifications and clearer clustering that reveal cellular heterogeneity and subtle subtypes. Its strong performance across datasets and batch conditions highlights robust generalization, making RegFormer a versatile tool for single-cell transcriptomics. In GRN construction, RegFormer generates high-resolution networks that align closely with known pathways and transcription factor activities, as exemplified by its accurate identification of key regulators such as GATA3 and SPI1. Despite these advances, challenges remain in scaling to larger datasets and addressing the dependence on GRN completeness. Future work will incorporate dynamic and context-specific regulatory networks and extend the framework to multi-omics integration for more comprehensive modeling of cellular states. Collectively, these results establish RegFormer as a powerful foundation model that unifies biological priors with advanced machine learning to enhance robustness, and discovery potential in single-cell research.

## Methods

### Data Collection and Preprocessing

We curated a large-scale corpus comprising 26 million human single cells across 45 tissues from the CZ CELLxGENE database, encompassing multiple sequencing platforms, developmental stages, and disease contexts. Each dataset was converted into an AnnData object and processed through a standardized pipeline to ensure consistency and scalability for model pretraining. As shown in Supplementary **Fig. S1A**, cells with fewer than 200 detected genes and genes outside the defined vocabulary were removed. Datasets were checked for scaling status, normalized using scanpy.pp.normalize_total when raw count data were detected, and log-transformed with scanpy.pp.log1p when the maximum expression value exceeded 25. Zero-expressed genes were filtered, and all processed AnnData objects were serialized into LMDB format to facilitate efficient data streaming during large-scale pretraining. Dataset distributions across tissues, sequencing platforms, developmental stages, and disease states are summarized in Supplementary **Fig. S1B-E**, highlighting the diversity and coverage of the pretraining corpus.

### Construction of Guided Gene Regulatory Networks

We constructed guided gene regulatory networks (GRNs) to encode transcriptional dependencies that inform model pretraining. As illustrated in **Supplementary Fig. S2A**, transcription factor (TF) motif-gene associations were derived from curated motif-scanning results (hg19-tss-centered-10kb-10species.mc9nr.genes_vs_motifs.rankings.feather) and mapped to TF identities using the motif annotation table (motifs-v9-nr.hgnc-m0.001-o0.0.tbl). Genes were filtered based on the predefined vocabulary to ensure consistency with the pretraining corpus.

For each TF, a directed edge was established to its top-20k ranked target genes, generating an initial regulatory graph G. Node embeddings were then learned using the Node2Vec algorithm, which captures higher-order connectivity patterns within the graph. To eliminate feedback loops and ensure acyclicity, edges were iteratively removed based on cosine similarity between node embeddings. In each cycle, the edge with the lowest similarity was deleted until a directed acyclic graph (DAG) was obtained. This process retained over 92% of edges while preserving degree and path-length distributions (**Supplementary Figs. S2B-F**). The final DAG was exported in DGL (.dgl) formats for downstream integration into the RegFormer pretraining pipeline.

## RegFormer Architecture

### Model Overview

RegFormer is a foundation model for single-cell transcriptomics that integrates gene regulatory network (GRN) priors with a state space-based sequence modeling framework. Each cell is represented as an ordered gene sequence derived from the GRN topology, allowing the model to capture both expression magnitude and regulatory context. The architecture consists of a dual-embedding input layer, a stack of Mamba-based state space blocks for contextual encoding, and multiple pretraining objectives for self-supervised representation learning.

### Topological Sorting

Input genes were ordered according to a directed acyclic gene regulatory network (GRN), denoted as, ***G*** = (***V***, ***E***)where nodes ***V*** represent genes and directed edges (*g_i_*→*g_j_*) in ***E*** denote transcriptional regulation from gene *g_i_* to gene *g_j_*.

To preserve causal directionality, nodes were arranged by topological depth, ensuring that upstream regulators always precede their downstream targets within each input sequence. Formally, the topological order

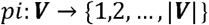

satisfies the condition

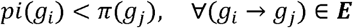

which guarantees that information propagates along biologically valid regulatory paths during sequence modeling.

In implementation, topological sorting was performed using the dgl.topological_nodes_generator function, which iteratively yields node batches from the source layer to the sink layer of the acyclic GRN, providing a depth-wise traversal consistent with the model’s sequential encoding.

For ablation experiments, randomized and edge-perturbed graphs ***G′ = (V, E′)*** were generated by shuffling or removing a subset of edges while maintaining node identity and approximate degree distribution.

### Dual-Embedding Scheme

Each gene token was represented by two complementary embeddings: a token embedding encoding gene identity and a value embedding encoding normalized expression magnitude. The two embeddings were summed and linearly projected into a shared latent space before entering the encoder. Formally, the representation of gene *g_i_* is given by

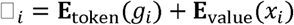

where **E**_token_ maps the discrete gene index *g_i_* to a learnable embedding vector, and **E**_value_ projects the normalized expression value *x_i_* through a linear transformation. The resulting sequence **H**_0_ = [**h**_1_, **h**_2_, …, **h**_*N*_] serves as input to the Mamba encoder stack. This dual scheme enables the model to jointly learn categorical and quantitative aspects of gene regulation and supports robust generalization across datasets with heterogeneous expression scales.

### Mamba-Based State Space Blocks

The core encoder of RegFormer consists of stacked Mamba blocks, each implementing a selective state space model (SSM) that efficiently captures long-range dependencies in gene sequences. The state space dynamics are defined as

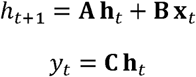

where **h***_t_* denotes the hidden state at position *t*, **x***_t_* the input embedding, and **A**, **B**, **C** are learnable transition, input, and output matrices, respectively.

In the Mamba implementation, these continuous dynamics are approximated through depthwise convolution and gated activation mechanisms, expressed as

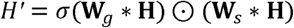

where * denotes a one-dimensional convolution along the sequence dimension, *σ* is a sigmoid gating function, and ʘ represents elementwise multiplication. This formulation enables selective state updatesc and efficient modeling of directional regulatory dependencies with linear computational complexity in sequence length.

Compared with attention-based architectures, the Mamba-based state space formulation allows RegFormer to capture global regulatory context with reduced memory usage and improved scalability across long gene sequences.

### Pretraining Objectives

RegFormer was pretrained using multiple self-supervised objectives designed to capture complementary aspects of transcriptional regulation. Three core tasks were implemented: Masked Value Prediction (MVP), Gene Expression Prediction for Cells (GEPC), and Regulatory Topology Modeling (TOPO). All objectives were jointly optimized under a weighted sum loss function.

To encourage local feature reconstruction, a subset of gene expression values was randomly masked, and the model was trained to predict their original magnitudes based on surrounding context. The loss was computed as mean squared error between predicted and observed values:

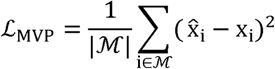

where □ denotes the set of masked positions, *x_i_* is the true normalized expression value, and 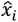 is the model’s prediction.

To align cell- and gene-level representations, a masked value decoder (MVC) was trained to reconstruct expression profiles from cell embeddings and query-specific gene vectors. For a given cell embedding ***c*** and gene query vectors **q***_i_*, the decoder computes

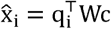

where **W** is a learnable projection. The resulting mean squared error loss encourages the cell embedding to encode global expression state:

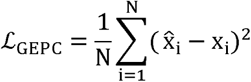

To learn regulatory directionality, the model was trained to predict the next gene token in a topologically sorted GRN sequence. Given the predicted logits **z***_t_* and target indices *y_t_*, a cross-entropy objective was applied:

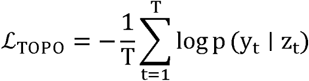

where masking was applied to non-regulatory or padding tokens to prevent trivial reconstruction. This objective enforces that embeddings respect the causal direction of transcriptional regulation.

All pretraining objectives were combined using adaptive weighting:

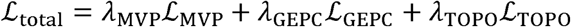

where λ*_i_* are task-specific coefficients chosen to balance gradient magnitudes. The masking ratio was set to 40% for all tasks, and loss terms were dynamically sanitized to prevent numerical instabilities during mixed-precision distributed training.

## Systematic Ablations

### Ablation Design

To dissect the contributions of individual architectural and pretraining components, we conducted a series of controlled ablation experiments (**Supplementary Figs. S4**). Each variant was initialized from the same configuration as the main RegFormer model and trained under identical optimization settings on **1 million blood cells** to ensure fair comparability. All experiments were performed using distributed data parallelism on 20 NVIDIA A100 GPUs (40 GB each).

To isolate the effects of core architectural modules, several variants were derived from the base Mamba framework. In the ***Random Sequence (Random)*** configuration, gene order within each input sequence was randomly permuted to remove graph-guided topological sorting. The ***Value-Only Embedding (OnlyValueEmb)*** variant excluded the gene identity token embedding, retaining only quantitative value embeddings to assess the contribution of gene-level contextual information. The ***Transformer Substitution (Transformer)*** variant replaced Mamba blocks with Transformer encoder layers of equal hidden dimension (256), feed-forward expansion (×4), and dropout rate (0.1), enabling a direct comparison between state-space and attention-based modeling. Finally, the ***Classification Head (BinCls)*** variant reformulated the value prediction task as a categorical classification problem optimized with a cross-entropy loss instead of mean squared error, testing sensitivity to prediction granularity.

To examine the influence of GRN topology on representation learning, two structural perturbations were introduced. In the ***Edge Shuffling (Shuffled)*** variant, 30% of edges were randomly reassigned while preserving node-degree distribution, thereby disrupting local connectivity but maintaining overall graph density. In the ***Edge Removal (Removed)*** variant, 30% of regulatory edges were randomly deleted to evaluate robustness under incomplete GRN priors. Both perturbations maintained consistent node identity and vocabulary alignment across models.

The effects of pretraining objectives were further examined by selectively removing or isolating self-supervised tasks. Three objectives—Masked Value Prediction (MVP), Gene Expression Prediction (GEPC), and Regulatory Role Prediction (TOPO)—were evaluated through ***Single-Objective variants (only MVP, only NTP, only GEPC)*** that retained one task and ***Objective-Removal variants (w/o MVP, w/o NTP, w/o GEPC)*** that excluded one task while keeping the remaining two active. In addition, a ***Masked Language Modeling (MLM)*** *variant* replaced continuous-value masking with discrete token-level masking, analogous to classical language modeling objectives in NLP, to assess how token-based context reconstruction affects representation learning. All ablation models used a consistent masking ratio (40%) and loss-balancing scheme identical to the main model.

To assess the effect of input coverage, two sequence lengths, 1.2k and 10k genes, were used across all ablation settings. Both configurations shared identical architectures, masking ratios, and optimization parameters.

Together, these systematically designed ablations established a unified framework for quantifying how regulatory priors, architectural components, and pretraining objectives collectively shape RegFormer’s learning behavior and representational capacity.

### GNN comparison

To assess the contribution of regulatory priors and state-space modeling, we compared RegFormer with graph neural network (GNN)–based pretraining frameworks operating on global gene co-expression graphs. The comparative GNN model adopted a four-layer relational graph attention network (Relational GATv2) with eight attention heads per layer, hidden dimension 512, ReLU activation, and dropout 0.2.

Global gene graphs were constructed by computing pairwise Pearson correlations among 3,000 highly variable genes and retaining the top 30 positively correlated partners (|r| > 0.2) per gene as edges, yielding an undirected weighted adjacency matrix used for all GNN training. The model was pretrained using two self-supervised objectives, masked-value reconstruction of gene expression and masked-edge prediction, optimized jointly with Adam under a cosine decay schedule (γ = 0.9), batch size 256, and automatic mixed precision (AMP). Approximately 15% of nodes and edges were randomly masked per batch.

Training proceeded for five epochs with early stopping based on validation loss. Node embeddings from the final layer were extracted for downstream analyses, including cell embedding, clustering, and GRN reconstruction. All models used identical data preprocessing, vocabulary, and evaluation settings as RegFormer to ensure comparability.

### Performance Assessment

To evaluate the representational quality of RegFormer, we performed complementary analyses at both the gene and cell levels. At the gene level, we examined whether the embeddings captured regulatory hierarchy through two tasks. In Task 1, each node in the gene regulatory network (GRN) was labeled as a transcription factor (TF) if it had outgoing edges, a target if it had only incoming edges, or isolated if unconnected. A logistic regression classifier was trained to distinguish TFs from non-TFs using 60/40 stratified splits, repeated 20 times with different random seeds, and mean macro-precision and macro-recall were reported. In Task 2, canonical correlation analysis (CCA) was performed between node embeddings and their in- and out-degree vectors to quantify the alignment between embedding geometry and regulatory connectivity. Higher classification accuracy and stronger correlations indicated that RegFormer embeddings more effectively encoded transcriptional directionality and hierarchical organization than randomized or ablated variants. At the cell level, embeddings were extracted from the final encoder layer, and the Adjusted Silhouette Width (ASW) for cell type was computed across **eight blood validation datasets** to assess biological separability and cluster coherence.

### Downstream Tasks

All downstream benchmarking of baseline single-cell foundation models was carried out within the BioLLM framework, which provides a standardized and reproducible pipeline encompassing data preprocessing, embedding generation, and task-specific evaluation using datasets listed in **Supplementary Table S1**.

### Cell Embedding

Cell embedding extraction and benchmarking of existing single-cell foundation models were performed using the BioLLM framework to maintain a consistent preprocessing and evaluation pipeline. Each model followed its native input configuration and feature constraints. Geneformer and scGPT, limited by maximum sequence lengths of 2,048 and 1,200 tokens, respectively, were provided with the top 3,000 highly variable genes (HVGs) as input features. In contrast, RegFormer, scBERT, and scFoundation processed complete gene expression matrices without gene filtering to retain full transcriptomic coverage.

Gene expression values were normalized according to each model’s original preprocessing protocol. RegFormer generated cell embeddings through three alternative pooling strategies, while scGPT used the CLS token representation, Geneformer averaged non-padded token embeddings, and scFoundation applied max pooling.

To evaluate the embedding quality, Average Silhouette Width (ASW) was calculated for both cell type and batch labels across multiple tissues. ASW was computed under two conditions, using all genes or 3,000 HVGs, to assess the effect of gene selection, and additional experiments varied the number of HVGs from 2,000 to 15,000. All analyses were implemented using the BioLLM framework, with Scanpy for data preprocessing, UMAP for visualization, and scikit-learn for ASW computation.

### Cell Type Annotation

Cell type annotation was performed in both intra-dataset and inter-dataset settings to evaluate each foundational model’s classification performance. Each dataset was divided into training and testing subsets at an 80:20 ratio, with the training portion further split into training and validation sets (90:10). Full gene expression profiles were retained to ensure consistency across models, and 3,000 highly variable genes (HVGs) were selected for training.

For RegFormer, fine-tuning was conducted using a linear classification head attached to the average-pooled cell embedding derived from Mamba encoder outputs. All models were trained for 20 epochs, with learning rates set to their recommended defaults: 0.0001 for scGPT and scFoundation, 0.001 for scBERT, 0.00005 for Geneformer, and 0.0001 for RegFormer. The effect of fine-tuning vs scratch training were further evaluated on the Zheng68k dataset.

### Gene Regulatory Network Construction

Gene Regulatory Networks (GRNs) were reconstructed from pretrained model embeddings to evaluate gene–gene relationships. Gene expression matrices (cells x genes) were first input into the RegFormer pretrained model to generate gene representations, where each gene was encoded as a high-dimensional embedding vector. Pairwise cosine similarity was then computed between all gene embeddings to quantify transcriptional association strength, resulting in a symmetric gene-gene similarity matrix. This matrix was interpreted as a weighted gene regulatory network, in which nodes represent genes and edge weights correspond to the embedding-derived similarity scores. Transcription factors (TFs) were used as potential regulators, and their top-20 target genes were identified based on cosine similarity values. The resulting networks were subsequently analyzed for topological properties and biological coherence, serving as the foundation for downstream clustering and functional enrichment analyses.

### Genetic Perturbation

To evaluate how well single-cell foundational models capture causal regulatory relationships, we conducted a gene perturbation prediction task using the Norman and Adamson datasets from the GEARS benchmark. Each dataset profiles single-cell transcriptomic responses following CRISPR-based gene perturbations, providing a suitable testbed for assessing generalization beyond observational data.

In this analysis, each foundational model provided pretrained gene embeddings obtained through representation learning across large-scale single-cell datasets. These embeddings were used to initialize the feature representations of genes within a co-expression-derived Genetic Relationship Graph, which serves as the structural backbone of the perturbation predictor. By replacing the default feature initialization with biologically informed embeddings, we aimed to examine whether pretrained regulatory representations enhance the model’s ability to infer transcriptional responses to perturbations.

All experiments were conducted under identical optimization settings to ensure fairness. The dimensionality of each embedding was aligned with the hidden size of the perturbation model, and training was performed for 20 epochs with a batch size of 32. Model parameters were fine-tuned while preserving the pretrained embedding space, allowing adaptation to perturbation-specific contexts without disrupting foundational gene representations. Performance was assessed on held-out perturbations by comparing predicted and observed gene expression changes.

### Drug Response

Drug response prediction was performed using the DeepCDR framework on two large pharmacogenomic resources: the Genomics of Drug Sensitivity in Cancer (GDSC), which provides half-maximal inhibitory concentration (IC□□) measurements across cancer cell lines, and the Cancer Cell Line Encyclopedia (CCLE), which contains matched gene expression, mutation, and DNA methylation profiles.

DeepCDR models drug efficacy by integrating chemical and molecular features through a multi-branch neural network. Drug molecular structures are represented as graphs and encoded by a graph convolutional network (GCN), while three parallel subnetworks independently extract latent biological features from CCLE mutation, expression, and methylation data. The outputs from all branches are fused through a convolutional aggregation module to predict IC□□ values for each drug–cell line pair.

To incorporate representations from single-cell foundational models, the CCLE gene expression profiles were first converted into embedding vectors using pretrained models such as RegFormer. Each cell line’s expression vector was passed through the pretrained model in inference mode to obtain a fixed-length embedding, which was normalized and used to replace the original expression input of DeepCDR’s transcriptomic subnetwork. This procedure allowed the model to leverage biologically structured representations learned from large-scale single-cell data.

For all experiments, the embedding dimension was matched to the hidden size of the DeepCDR architecture to ensure consistent input dimensionality. Training and testing were conducted on GPU devices under a leave-one-drug-out cross-validation protocol, in which data associated with a single drug were held out during training and used exclusively for evaluation. Each model was trained using the Adam optimizer with a batch size of 64 and learning rate of 1×10□□ for 50 epochs, and the best checkpoint was selected based on validation performance. Predictive accuracy was quantified by the Pearson correlation coefficient (PCC) and Spearman rank correlation between the predicted and observed ICLL values across test drug–cell pairs.

## Evaluation Metrics

### Cell Embedding Metrics

To assess the quality of learned cell embeddings, we evaluated both cell type clustering and batch effect removal within the embedding space. The primary evaluation metric was the Average Silhouette Width (ASW), which quantifies the balance between intra-cluster cohesion and inter-cluster separation.

For cell type clustering, ASW values range from 0 to 1, where 0 indicates poorly defined clusters, 0.5 suggests moderate overlap, and 1 reflects well-separated cell types. Higher ASW values indicate better-defined cell clusters, demonstrating more accurate embedding performance. Formally, the ASW for cell types is given by:

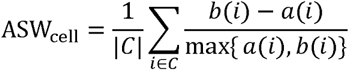

where *a*(*i*) is the mean intra-cluster distance of cell *i*, *b*(*i*), is the smallest mean distance to another cluster, and *C* is the set of all cell type labels.

For batch effect evaluation, ASW was computed using batch labels instead of cell types. A score of 0 indicates optimal mixing across batches, while deviations from 0 indicate batch effects. To facilitate comparison, batch ASW values were converted by subtracting them from 1, yielding:

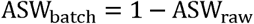

Thus, a score of 1 represents complete batch integration and 0 denotes total separation.

### Cell Type Annotation Metrics

The performance of cell type annotation was evaluated by comparing predicted and true labels using three complementary metrics: Accuracy, Macro-F1, and weighted average precision (Weighted AP). These metrics jointly assess both overall correctness and per-class balance, providing a robust evaluation of annotation performance across imbalanced cell populations.

The accuracy was defined as the proportion of correctly predicted labels among all cells:

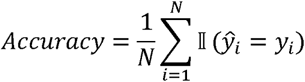

where *N* is the total number of cells, *y_i_*, is the true label, 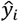 is the predicted label, and ∏ is the indicator function.

To account for class imbalance, the Macro-F1 score was computed as the average of per-class F1 scores, where each class-specific F1 is defined by the harmonic mean of its precision and recall:

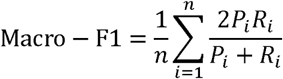

where *P_i_* and *R_i_* denote the precision and recall for class *i*, respectively.

In addition, to emphasize class frequency during evaluation, we computed the Weighted Average Precision (Weighted AP), which weights each class-specific average precision *AP_i_* by its relative cell proportion *w_i_*

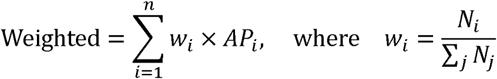

Here, *AP_i_* represents the area under the precision–recall curve for class *i*, and N*_i_* is the number of cells belonging to class *i* Weighted AP therefore reflects both prediction quality and dataset composition, offering a comprehensive measure of annotation performance.

### Gene Regulatory Network Metrics

To assess the biological coherence of model-derived Gene Regulatory Networks (GRNs), we computed a Functional Similarity Index (FSI) based on Gene Ontology (GO) Biological Process annotations. Genes sharing at least one GO term were connected to form a reference adjacency matrix *A*_GO_, while each model provided a corresponding gene–gene network *A*_model_ derived from embedding proximity. The FSI quantifies the agreement between model-inferred and GO-based connectivity using cosine similarity:

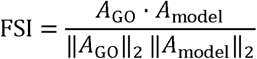

Higher values indicate stronger alignment with functional gene relationships.

The k-means clustering algorithm was applied to the model-derived gene networks to identify gene modules based on their connectivity patterns. By varying the number of clusters from 5 to 30 in increments of 5, we explored the modular structure of reconstructed GRNs at different granularities. For each clustering level, Gene Ontology (GO) enrichment analysis was performed on the resulting modules, and the number of significantly enriched Biological Process terms *P*_adj_ < 0.05 was recorded. Models with higher enrichment counts demonstrated stronger biological coherence, as illustrated in the GO (Biological Process) enrichment curves across clustering resolutions.

### Genetic Perturbation Metrics

To benchmark models on perturbation response prediction, we followed the GEARS evaluation protocol. Performance was assessed using the Mean Squared Error (MSE) and Pearson Correlation Coefficient (PCC) between predicted and experimentally observed differential gene expression values:

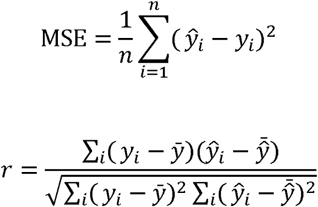

where *y_i_* and 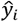 denote the observed and predicted differential expressions, respectively.

### Drug Response Metrics

For drug sensitivity prediction, we used preprocessed data from DeepCDR and evaluated model performance using both the Pearson Correlation Coefficient (PCC) and the Spearman’s Rank Correlation Coefficient (SRCC) between predicted and actual IC50 values:

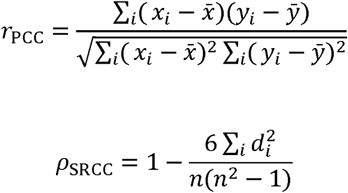

where *d_i_* is the rank difference between predicted and observed IC50 values for the *i*-th drug--cell pair. Higher PCC and SRCC values indicate stronger agreement between model predictions and experimental outcomes.

## Data availability

All benchmarking dataseting used in the research were collected from published sources, with detailed descriptions provided in **Supplementary Table S1**.

## Code availability

RegFormer is packaged, and distributed as an open-source, publicly available repository at https://github.com/BGIResearch/RegFormer.

## Acknowledgement

We acknowledge the Stomics Cloud platform (https://cloud.stomics.tech/) for providing GPU computational resources. We thank the colleagues in our research group for inspiring discussion and their contributions.

## Author contributions

Y.Z., Y.L. and S.F. conceptualized the study. L.H. was responsible for the model design and tool implementation. L.H., H.Q. Y.Z., Y.L. P.Q., and Z.G. performed data analysis and model evaluation. L.C., W.J., Q.C., Y.S., T.X. and Z.D. contributed key ideas and advice. L.H. wrote the manuscript. X.X, Y.L., S.F. and Y.Z. supervised the study.

