## Supplementary Figures for "RegFormer: A Single-Cell Foundation Model Powered by Gene Regulatory Hierarchies"

### Figure S1

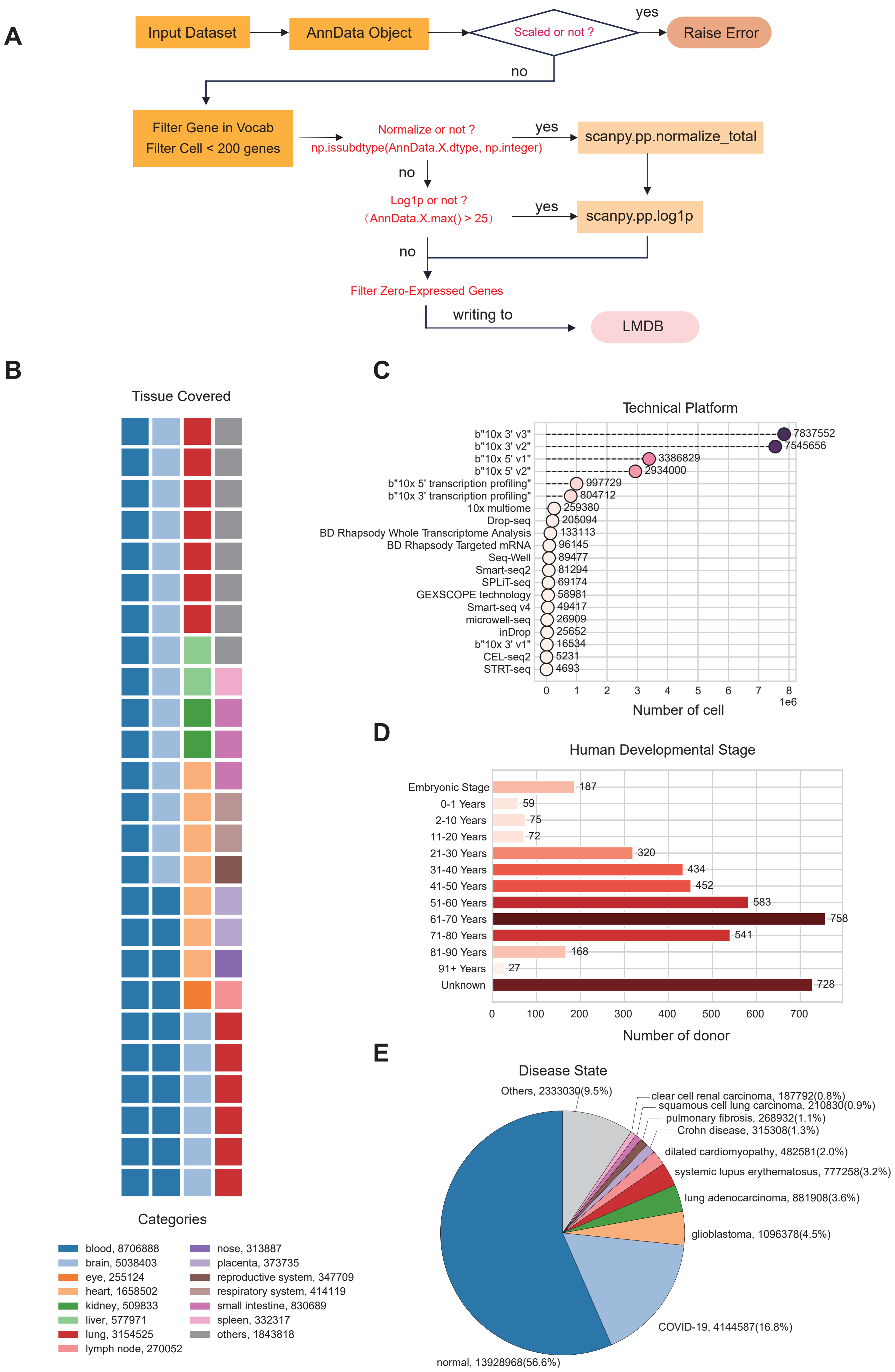

A
Input Dataset
AnnData Object
Scaled or not ?
yes
Raise Error
no
Filter Gene in Vocab
Filter Cell < 200 genes
Normalize or not ?
np.issubdtype(AnnData.X.dtype, np.integer)
yes
scanpy.pp.normalize_total
no
Log1p or not ?
（AnnData.X.max() > 25）
yes
scanpy.pp.log1p
no
Filter Zero-Expressed Genes
writing to
LMDB
B
Tissue Covered
Categories
blood, 8706888
brain, 5038403
eye, 255124
heart, 1658502
kidney, 509833
liver, 577971
lung, 3154525
lymph node, 270052
nose, 313887
placenta, 373735
reproductive system, 347709
respiratory system, 414119
small intestine, 830689
spleen, 332317
others, 1843818
C
Technical Platform
b"10x 3' v3" 7837552
b"10x 3' v2" 7545656
b"10x 5' v1" 3386829
b"10x 5' v2" 2934000
b"10x 5' transcription profiling" 997729
b"10x 3' transcription profiling" 804712
10x multiome 259380
Drop-seq 205094
BD Rhapsody Whole Transcriptome Analysis 133113
BD Rhapsody Targeted mRNA 96145
Seq-Well 89477
Smart-seq2 81294
SPLiT-seq 69174
GEXSCOPE technology 58981
Smart-seq v4 49417
microwell-seq 26909
inDrop 25652
b"10x 3' v1" 16534
CEL-seq2 5231
STRT-seq 4693
0 1 2 3 4 5 6 7 8
1e6
Number of cell
D
Human Developmental Stage
Embryonic Stage 187
0-1 Years 59
2-10 Years 75
11-20 Years 72
21-30 Years 320
31-40 Years 434
41-50 Years 452
51-60 Years 583
61-70 Years 758
71-80 Years 541
81-90 Years 168
91+ Years 27
Unknown 728
0 100 200 300 400 500 600 700
Number of donor
E
Disease State
Others, 2333030(9.5%)
clear cell renal carcinoma, 187792(0.8%)
squamous cell lung carcinoma, 210830(0.9%)
pulmonary fibrosis, 268932(1.1%)
Crohn disease, 315308(1.3%)
dilated cardiomyopathy, 482581(2.0%)
systemic lupus erythematosus, 777258(3.2%)
lung adenocarcinoma, 881908(3.6%)
glioblastoma, 1096378(4.5%)
COVID-19, 4144587(16.8%)
normal, 13928968(56.6%)

**Figure S1. Data preprocessing and overview of the pretraining dataset.**

**(A)** Workflow of data preprocessing and LMDB construction.

**(B)** Tissue types covered in the pretraining dataset.

**(C)** Summary of technical platforms used for data generation.

**(D)** Distribution of donor samples across human developmental stages.

**(E)** Proportion of samples across different disease states.

### Figure S2

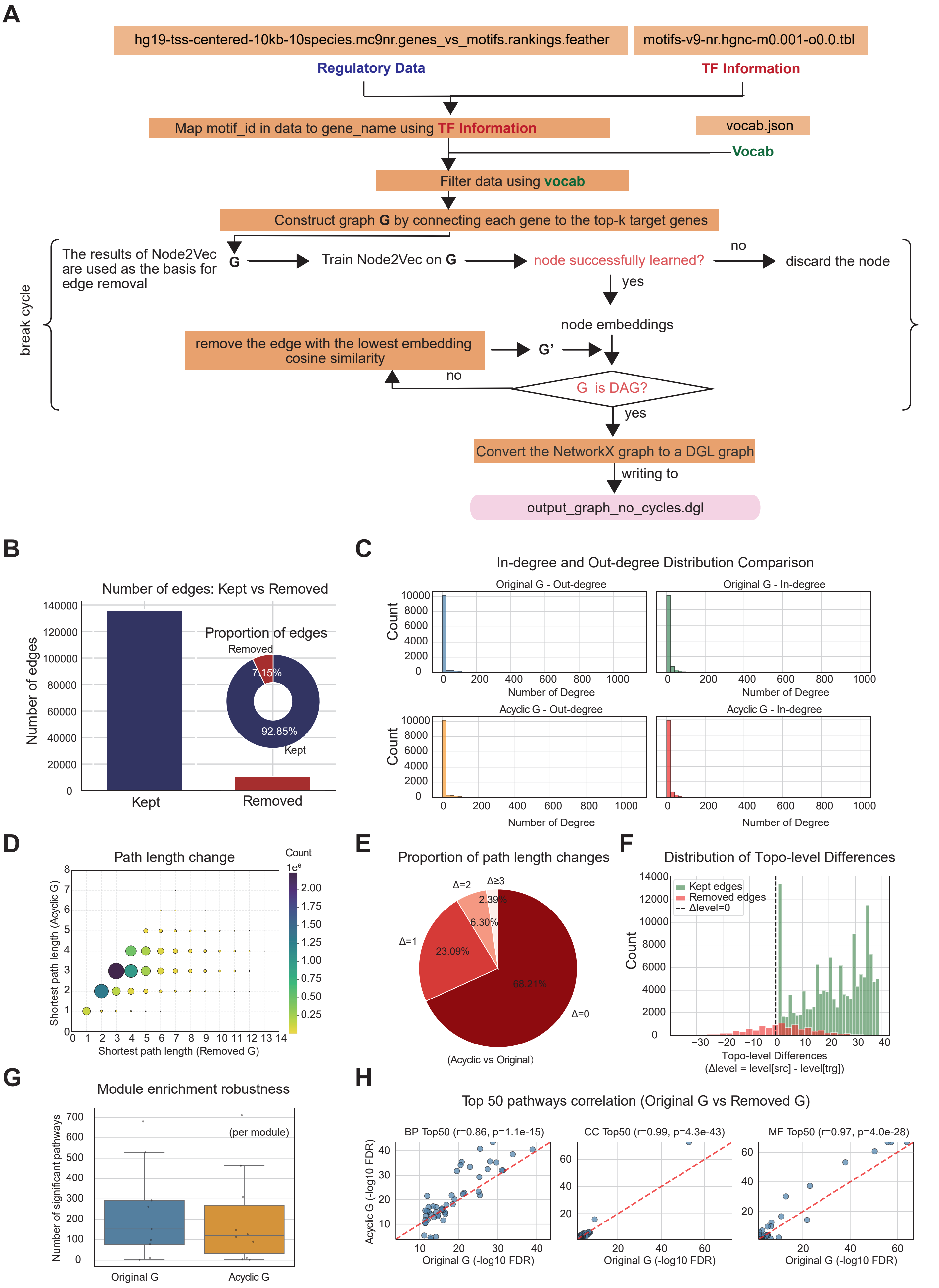

A
hg19-tss-centered-10kb-10species.mc9nr.genes_vs_motifs.rankings.feather
motifs-v9-nr.hgnc-m0.001-o0.0.tbl
Regulatory Data
TF Information
Map motif_id in data to gene_name using TF Information
vocab.json
Vocab
Filter data using vocab
Construct graph G by connecting each gene to the top-k target genes
break cycle
The results of Node2Vec are used as the basis for edge removal
G
Train Node2Vec on G
node successfully learned?
no
discard the node
yes
node embeddings
remove the edge with the lowest embedding cosine similarity
G'
no
G is DAG?
yes
Convert the NetworkX graph to a DGL graph
writing to
output_graph_no_cycles.dgl
B
Number of edges: Kept vs Removed
Number of edges
Proportion of edges
Removed
7.15%
92.85%
Kept
Kept
Removed
C
In-degree and Out-degree Distribution Comparison
Original G - Out-degree
Original G - In-degree
Acyclic G - Out-degree
Acyclic G - In-degree
Count
Number of Degree
D
Path length change
Count
1e6
Shortest path length (Acyclic G)
Shortest path length (Removed G)
E
Proportion of path length changes
Δ=2
Δ≥3
2.39%
6.30%
Δ=1
23.09%
68.21%
Δ=0
(Acyclic vs Original)
F
Distribution of Topo-level Differences
Kept edges
Removed edges
Δlevel=0
Count
Topo-level Differences
(Δlevel = level[src] - level[trg])
G
Module enrichment robustness
Number of significant pathways
(per module)
Original G
Acyclic G
H
Top 50 pathways correlation (Original G vs Removed G)
BP Top50 (r=0.86, p=1.1e-15)
CC Top50 (r=0.99, p=4.3e-43)
MF Top50 (r=0.97, p=4.0e-28)
Acyclic G (-log10 FDR)
Original G (-log10 FDR)

**Figure S2. Construction and evaluation of the GRN graph.**

**(A)** Workflow of acyclic GRN graph construction based on motif – gene ranking data, TF information, and gene vocab.

**(B)** Proportion of edges retained and removed during cycle removal.

**(C)** Comparison of in-degree and out-degree distributions before and after cycle removal.

**(D)** Changes in shortest path lengths between the original and acyclic graphs.

**(E)** Proportion of path length changes between the original and acyclic graphs.

**(F)** Distribution of topological level differences between the original and acyclic graphs.

**(G)** Comparison of module enrichment robustness before and after cycle removal.

**(H)** Correlation of the top 50 enriched pathways between the original and acyclic graphs.

### Figure S3

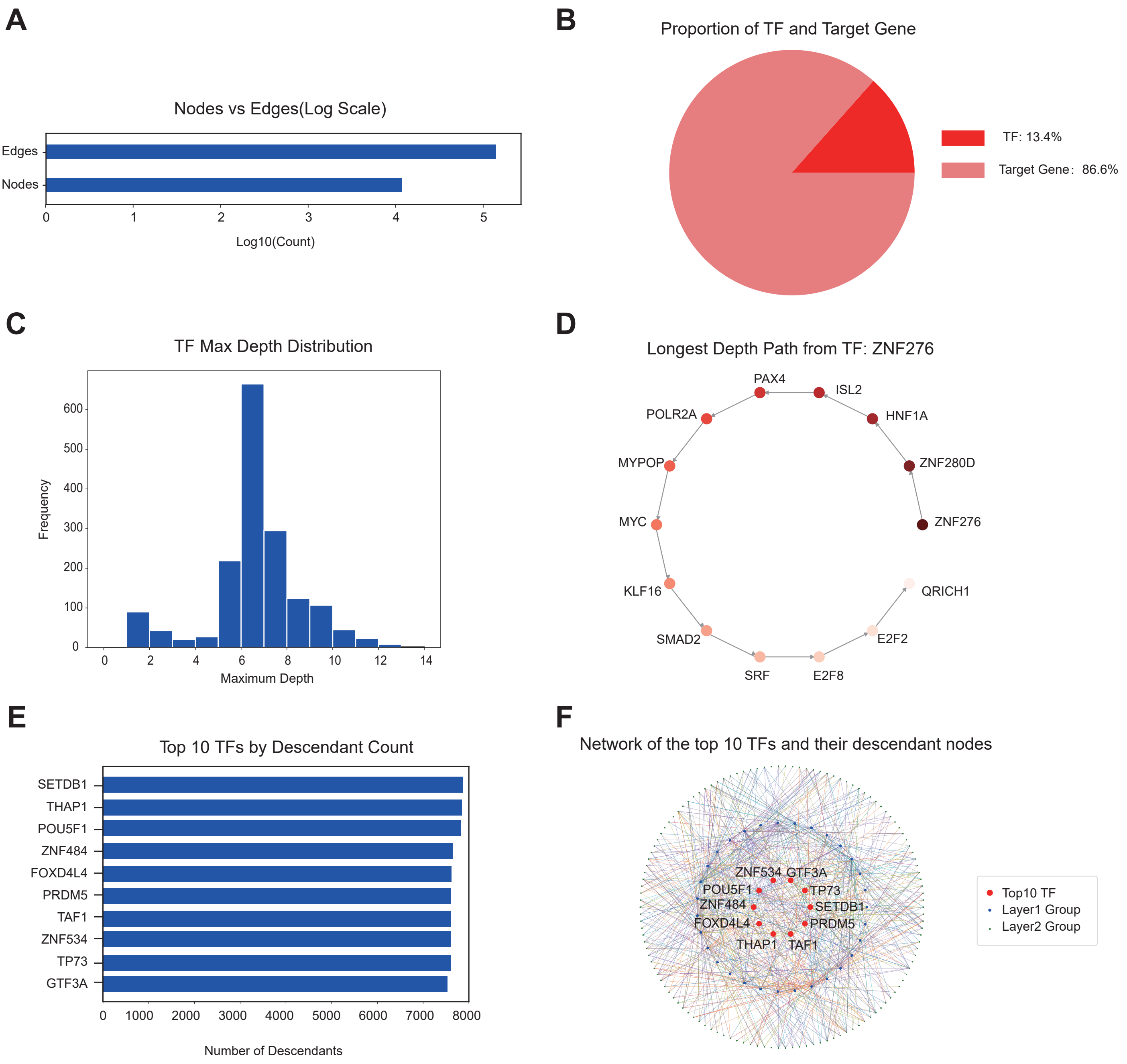


**Figure S3. Basic description of the acyclic GRN graph.**

**(A)** Comparison of node and edge counts on a logarithmic scale.

**(B)** Proportion of transcription factors (TFs) and target genes in the acyclic graph.

**(C)** Distribution of the maximum regulatory depth among transcription factors.

**(D)** Example of the longest regulatory path originating from TF ZNF276.

**(E)** Top 10 transcription factors ranked by the number of descendant genes.

**(F)** Network visualization of the top 10 transcription factors and their descendant nodes organized by hierarchical layers.

### Figure S4

| Experiment | Main | Random | OnlyValueEmb | Shuffled | Removed | Transformer | BinCls | MLM | w/o MVP | w/o NTP | w/o GEPC | only MVP | only NTP | only GEPC |
|---|---|---|---|---|---|---|---|---|---|---|---|---|---|---|
| **Description** | Full model | Randomly Sorted Sequence | Keep only value embedding | 30% edges shuffled | 30% edges removed | Transformer architecture | Value prediction as classification | Gene prediction with MLM | Remove MVP task | Remove NTP task | Remove GEPC task | Use only MVP task | Use only NTP task | Use only GEPC task |
| **Archetecture** | Mamba | Mamba | Mamba | Mamba | Mamba | **Transformer** | Mamba | Mamba | Mamba | Mamba | Mamba | Mamba | Mamba | Mamba |
| **Graph** | Original | Original | Original | **Shuffled** | **Removed** | Original | Original | Original | Original | Original | Original | Original | Original | Original |
| **MVP** | ✓ | ✓ | ✓ | ✓ | ✓ | ✓ | ✓ | ✓ | ✗ | ✓ | ✓ | ✓ | ✓ | ✓ |
| **TOPO** | ✓ | ✓ | ✓ | ✓ | ✓ | ✓ | ✓ | ✓ | ✓ | ✗ | ✓ | ✗ | ✗ | ✓ |
| **GEPC** | ✓ | ✓ | ✓ | ✓ | ✓ | ✓ | ✓ | ✓ | ✓ | ✓ | ✗ | ✗ | ✓ | ✗ |
| **Token Prediction** | NTP | NTP | NTP | NTP | NTP | NTP | NTP | **MLM** | NTP | NTP | NTP | NTP | NTP | NTP |
| **Value Prediction** | MSE | MSE | MSE | MSE | MSE | MSE | **CE** | MSE | MSE | MSE | MSE | MSE | MSE | MSE |
| **TOPO SORT** | ✓ | ✗ | ✓ | ✓ | ✓ | ✓ | ✓ | ✓ | ✓ | ✓ | ✓ | ✓ | ✓ | ✓ |
| **Dual Emb** | ✗ | ✗ | ✓ | ✗ | ✗ | ✗ | ✗ | ✗ | ✗ | ✗ | ✗ | ✗ | ✗ | ✗ |
| **1.2k** | ★★★★☆ | ★★★★☆ | ★⯪☆☆☆ | ★★★★☆ | ★★★★☆ | ★★★☆☆ | ★★★★☆ | ★★★★☆ | ★★★⯪☆ | ★★★★☆ | ★★★⯪☆ | ★★★★☆ | ★☆☆☆☆ | ★★★☆☆ |
| **10k** | ★★★★★ | ★★★★⯪ | ★★☆☆☆ | ★★★★⯪ | ★★★★★ | ★★★★⯪ | ★★★★★ | ★★★★⯪ | | | | | | |

**Figure S4. Ablation study of RegFormer across different training settings..**
Comparison of model variants with changes in architecture, graph structure, and pretraining objectives, showing their effects on ASW scores for sequence lengths of 1.2k and 10k.

### Figure S5

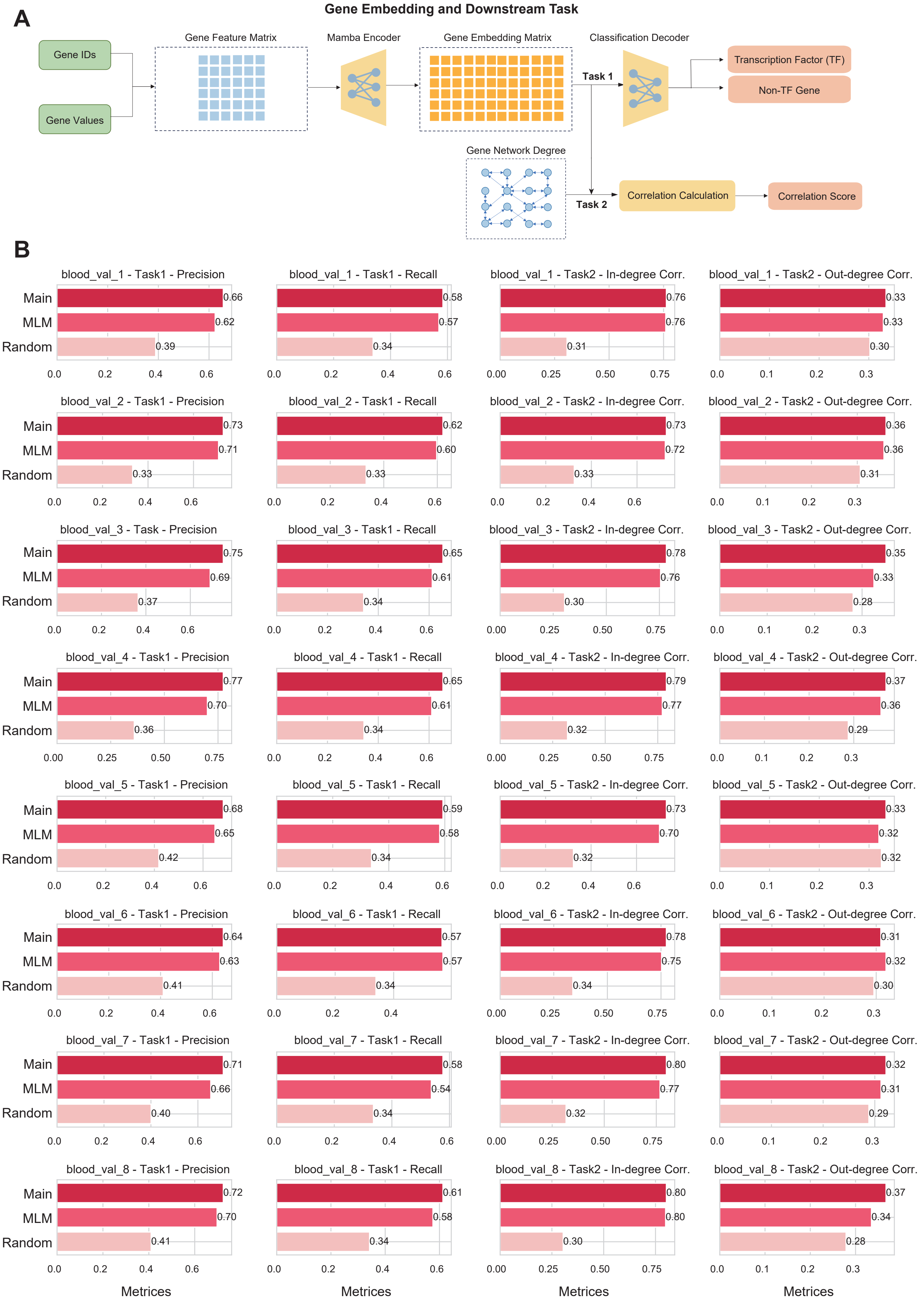

A
Gene Embedding and Downstream Task
Gene IDs
Gene Values
Gene Feature Matrix
Mamba Encoder
Gene Embedding Matrix
Classification Decoder
Task 1
Transcription Factor (TF)
Non-TF Gene
Gene Network Degree
Task 2
Correlation Calculation
Correlation Score
B
blood_val_1 - Task1 - Precision
Main 0.66
MLM 0.62
Random 0.39
blood_val_1 - Task1 - Recall
0.58
0.57
0.34
blood_val_1 - Task2 - In-degree Corr.
0.76
0.76
0.31
blood_val_1 - Task2 - Out-degree Corr.
0.33
0.33
0.30
blood_val_2 - Task1 - Precision
Main 0.73
MLM 0.71
Random 0.33
blood_val_2 - Task1 - Recall
0.62
0.60
0.33
blood_val_2 - Task2 - In-degree Corr.
0.73
0.72
0.33
blood_val_2 - Task2 - Out-degree Corr.
0.36
0.36
0.31
blood_val_3 - Task - Precision
Main 0.75
MLM 0.69
Random 0.37
blood_val_3 - Task1 - Recall
0.65
0.61
0.34
blood_val_3 - Task2 - In-degree Corr.
0.78
0.76
0.30
blood_val_3 - Task2 - Out-degree Corr.
0.35
0.33
0.28
blood_val_4 - Task1 - Precision
Main 0.77
MLM 0.70
Random 0.36
blood_val_4 - Task1 - Recall
0.65
0.61
0.34
blood_val_4 - Task2 - In-degree Corr.
0.79
0.77
0.32
blood_val_4 - Task2 - Out-degree Corr.
0.37
0.36
0.29
blood_val_5 - Task1 - Precision
Main 0.68
MLM 0.65
Random 0.42
blood_val_5 - Task1 - Recall
0.59
0.58
0.34
blood_val_5 - Task2 - In-degree Corr.
0.73
0.70
0.32
blood_val_5 - Task2 - Out-degree Corr.
0.33
0.32
0.32
blood_val_6 - Task1 - Precision
Main 0.64
MLM 0.63
Random 0.41
blood_val_6 - Task1 - Recall
0.57
0.57
0.34
blood_val_6 - Task2 - In-degree Corr.
0.78
0.75
0.34
blood_val_6 - Task2 - Out-degree Corr.
0.31
0.32
0.30
blood_val_7 - Task1 - Precision
Main 0.71
MLM 0.66
Random 0.40
blood_val_7 - Task1 - Recall
0.58
0.54
0.34
blood_val_7 - Task2 - In-degree Corr.
0.80
0.77
0.32
blood_val_7 - Task2 - Out-degree Corr.
0.32
0.31
0.29
blood_val_8 - Task1 - Precision
Main 0.72
MLM 0.70
Random 0.41
blood_val_8 - Task1 - Recall
0.61
0.58
0.34
blood_val_8 - Task2 - In-degree Corr.
0.80
0.80
0.30
blood_val_8 - Task2 - Out-degree Corr.
0.37
0.34
0.28
Metrices
Metrices
Metrices
Metrices

**Figure S5. Evaluation of gene embeddings in downstream tasks.**

**(A)** Workflow of gene embedding generation and downstream evaluation using classification and correlation tasks.

**(B)** Comparison of RegFormer (Main), MLM, and Random models across eight blood validation sets, showing performance in transcription factor classification (Task 1) and correlation with gene network degrees (Task 2).

### Figure S6

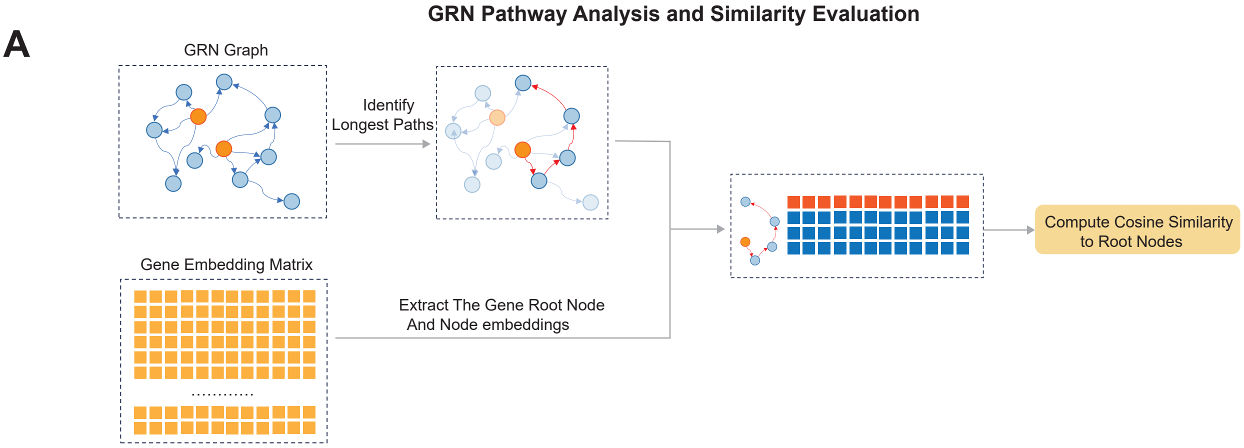
GRN Pathway Analysis and Similarity Evaluation
A
GRN Graph
Identify Longest Paths
Gene Embedding Matrix
Extract The Gene Root Node And Node embeddings
Compute Cosine Similarity to Root Nodes

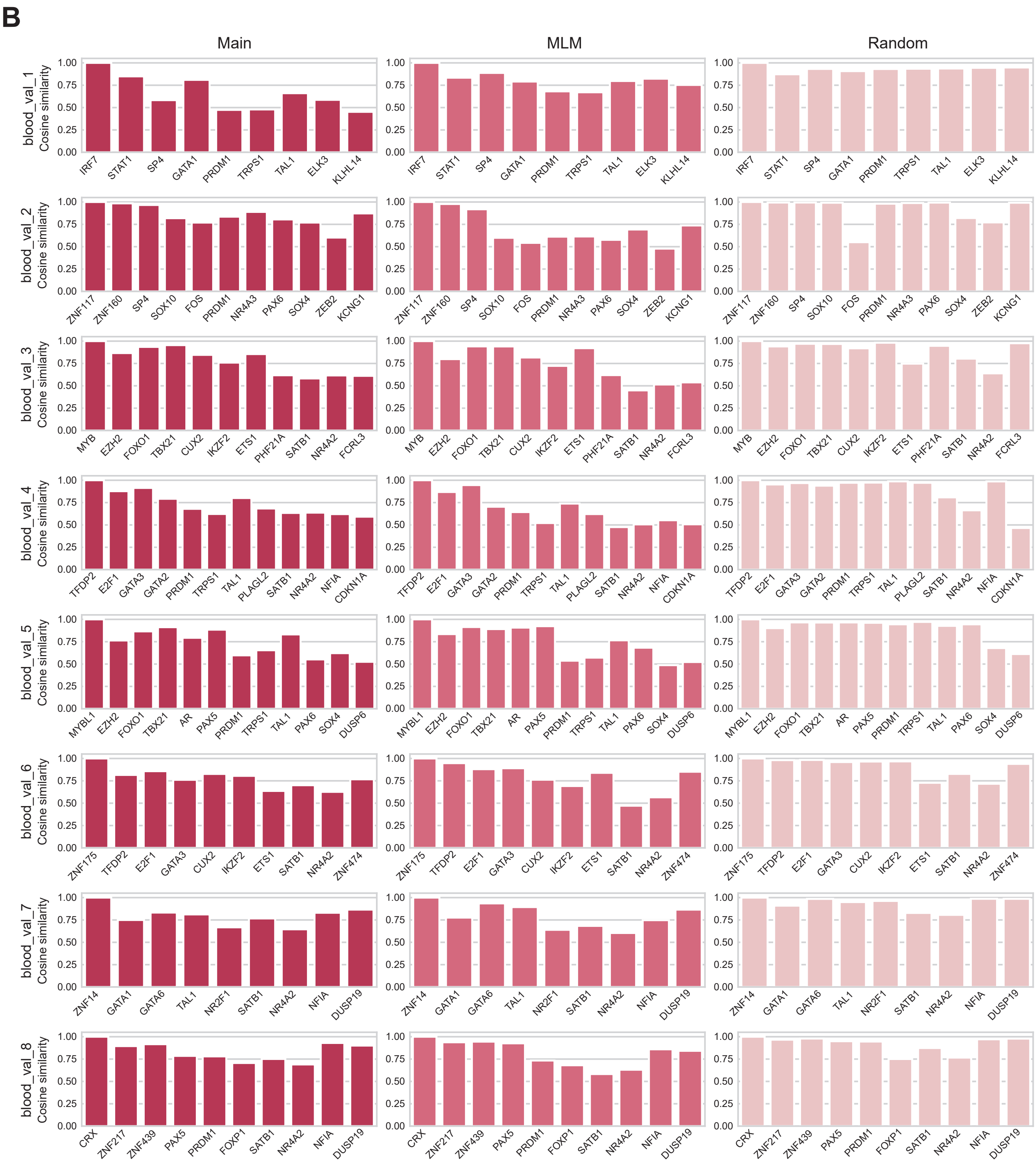
B
Main
MLM
Random
Cosine similarity
blood_val_1
IRF7 STAT1 SP4 GATA1 PRDM1 TRPS1 TAL1 ELK3 KLHL14
blood_val_2
ZNF117 ZNF160 SP4 SOX10 FOS PRDM1 NR4A3 PAX6 SOX4 ZEB2 KCNG1
blood_val_3
MYB EZH2 FOXO1 TBX21 CUX2 IKZF2 ETS1 PHF21A SATB1 NR4A2 FCRL3
blood_val_4
TFDP2 E2F1 GATA3 GATA2 PRDM1 TRPS1 TAL1 PLAGL2 SATB1 NR4A2 NFIA CDKN1A
blood_val_5
MYBL1 EZH2 FOXO1 TBX21 AR PAX5 PRDM1 TRPS1 TAL1 PAX6 SOX4 DUSP6
blood_val_6
ZNF175 TFDP2 E2F1 GATA3 CUX2 IKZF2 ETS1 SATB1 NR4A2 ZNF474
blood_val_7
ZNF14 GATA1 GATA6 TAL1 NR2F1 SATB1 NR4A2 NFIA DUSP19
blood_val_8
CRX ZNF217 ZNF439 PAX5 PRDM1 FOXP1 SATB1 NR4A2 NFIA DUSP19
0.00 0.25 0.50 0.75 1.00

**Figure S6. GRN pathway analysis and similarity evaluation using RegFormer embeddings.**
**(A)** Workflow for identifying the longest regulatory paths in the GRN and evaluating cosine similarity between root nosde embeddings and their descendant nodes.
**(B)** Comparison of RegFormer (Main), MLM, and Random models across eight blood validation sets, showing cosine similarity distributions of gene embeddings along GRN pathways.

### Figure S7

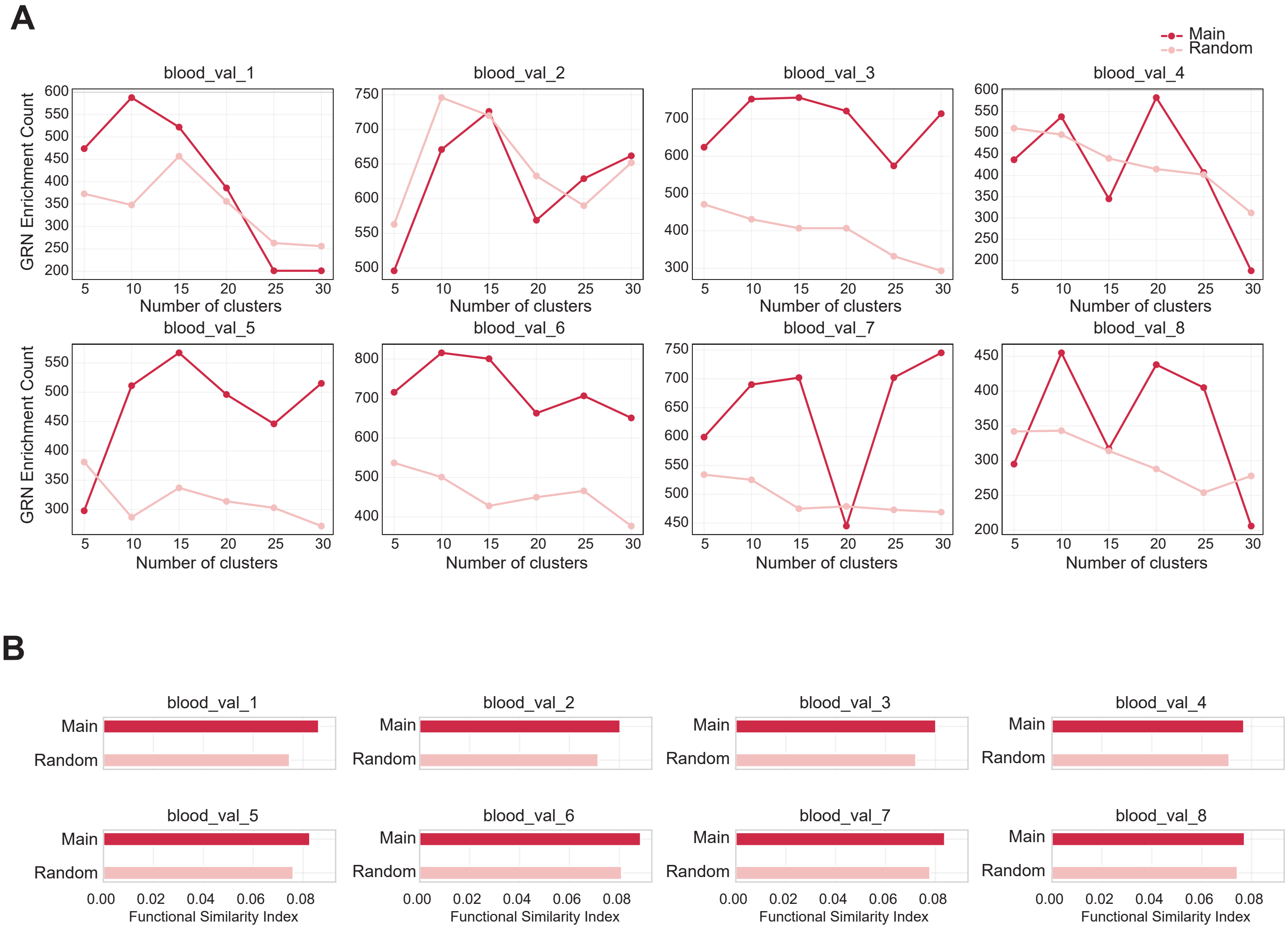


**Figure S7. Functional evaluation on reconstructed GRNs.**

**(A)** GRN enrichment counts across different clustering resolutions for eight blood validation datasets based on reconstructed GRNs.

**(B)** Functional similarity index across datasets comparing GRN-guided (Main) and non-GRN-guided (Random) models.

### Figure S8

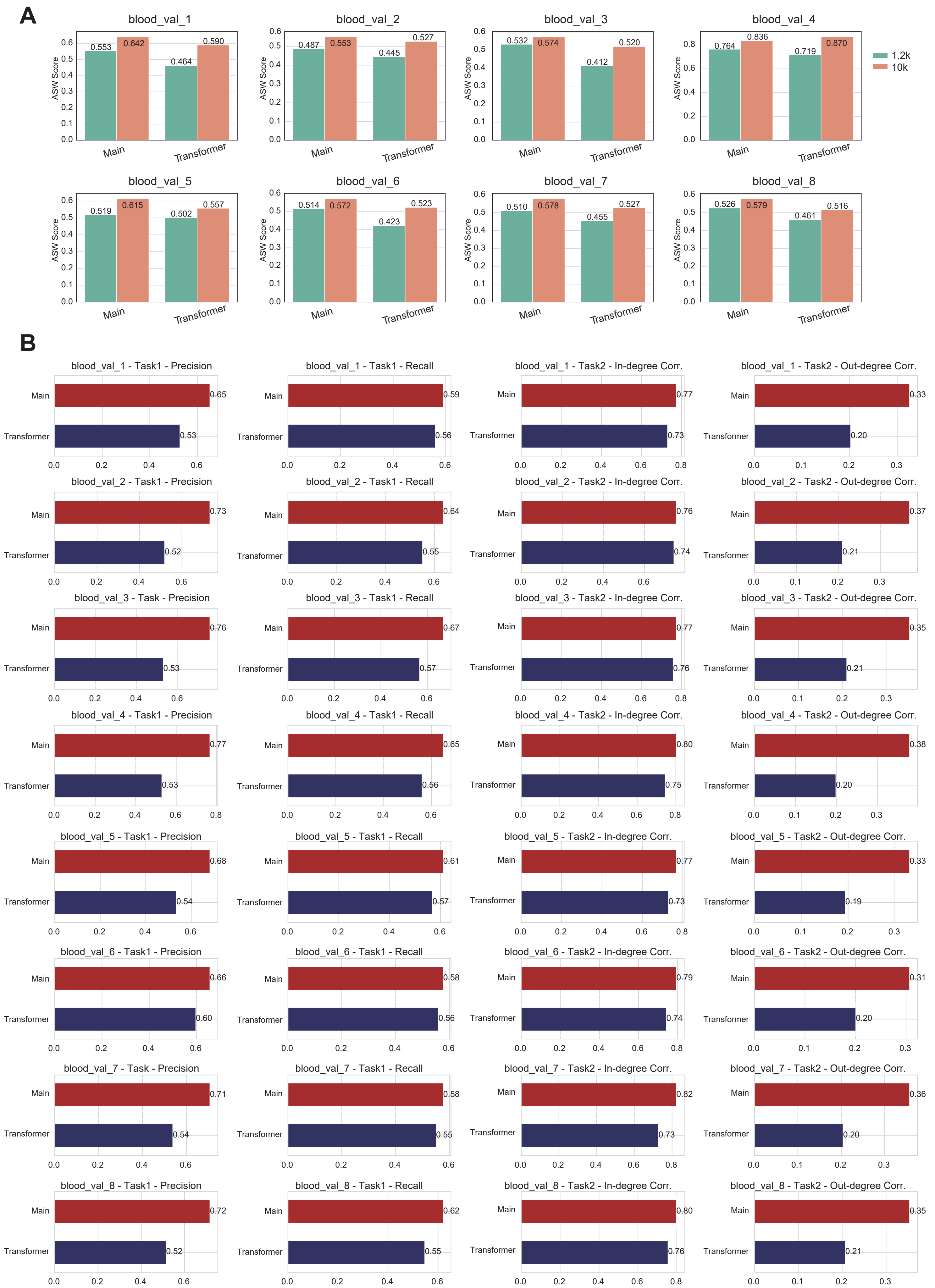
A
blood_val_1
ASW Score
0.553
0.642
0.464
0.590
Main
Transformer
blood_val_2
0.487
0.553
0.445
0.527
blood_val_3
0.532
0.574
0.412
0.520
blood_val_4
0.764
0.836
0.719
0.870
1.2k
10k
blood_val_5
0.519
0.615
0.502
0.557
blood_val_6
0.514
0.572
0.423
0.523
blood_val_7
0.510
0.578
0.455
0.527
blood_val_8
0.526
0.579
0.461
0.516
B
blood_val_1 - Task1 - Precision
Main 0.65
Transformer 0.53
blood_val_1 - Task1 - Recall
Main 0.59
Transformer 0.56
blood_val_1 - Task2 - In-degree Corr.
Main 0.77
Transformer 0.73
blood_val_1 - Task2 - Out-degree Corr.
Main 0.33
Transformer 0.20
blood_val_2 - Task1 - Precision
Main 0.73
Transformer 0.52
blood_val_2 - Task1 - Recall
Main 0.64
Transformer 0.55
blood_val_2 - Task2 - In-degree Corr.
Main 0.76
Transformer 0.74
blood_val_2 - Task2 - Out-degree Corr.
Main 0.37
Transformer 0.21
blood_val_3 - Task - Precision
Main 0.76
Transformer 0.53
blood_val_3 - Task1 - Recall
Main 0.67
Transformer 0.57
blood_val_3 - Task2 - In-degree Corr.
Main 0.77
Transformer 0.76
blood_val_3 - Task2 - Out-degree Corr.
Main 0.35
Transformer 0.21
blood_val_4 - Task1 - Precision
Main 0.77
Transformer 0.53
blood_val_4 - Task1 - Recall
Main 0.65
Transformer 0.56
blood_val_4 - Task2 - In-degree Corr.
Main 0.80
Transformer 0.75
blood_val_4 - Task2 - Out-degree Corr.
Main 0.38
Transformer 0.20
blood_val_5 - Task1 - Precision
Main 0.68
Transformer 0.54
blood_val_5 - Task1 - Recall
Main 0.61
Transformer 0.57
blood_val_5 - Task2 - In-degree Corr.
Main 0.77
Transformer 0.73
blood_val_5 - Task2 - Out-degree Corr.
Main 0.33
Transformer 0.19
blood_val_6 - Task1 - Precision
Main 0.66
Transformer 0.60
blood_val_6 - Task1 - Recall
Main 0.58
Transformer 0.56
blood_val_6 - Task2 - In-degree Corr.
Main 0.79
Transformer 0.74
blood_val_6 - Task2 - Out-degree Corr.
Main 0.31
Transformer 0.20
blood_val_7 - Task - Precision
Main 0.71
Transformer 0.54
blood_val_7 - Task1 - Recall
Main 0.58
Transformer 0.55
blood_val_7 - Task2 - In-degree Corr.
Main 0.82
Transformer 0.73
blood_val_7 - Task2 - Out-degree Corr.
Main 0.36
Transformer 0.20
blood_val_8 - Task1 - Precision
Main 0.72
Transformer 0.52
blood_val_8 - Task1 - Recall
Main 0.62
Transformer 0.55
blood_val_8 - Task2 - In-degree Corr.
Main 0.80
Transformer 0.76
blood_val_8 - Task2 - Out-degree Corr.
Main 0.35
Transformer 0.21

**Figure S8. Comparison between Mamba-based and Transformer-based architectures.**

**(A)** ASW scores across eight blood validation datasets for RegFormer (Main) and Transformer variants at sequence lengths of 1.2k and 10k.

**(B)** Performance comparison on gene-level downstream tasks, including Task 1 (Precision, Recall) and Task 2 (In-degree and Out-degree Correlation), across validation datasets.

### Figure S9

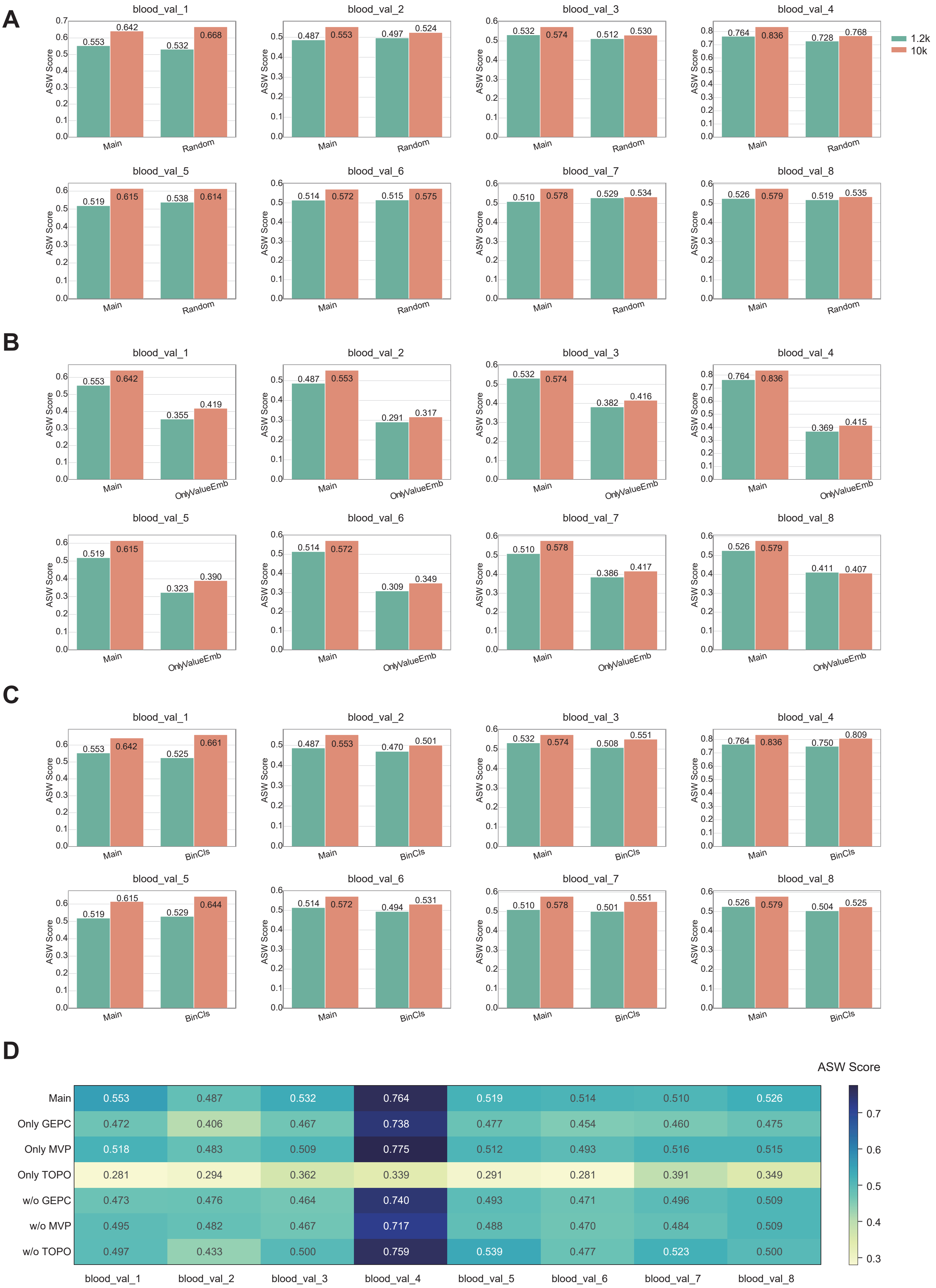
A
blood_val_1
0.553 0.642 0.532 0.668
blood_val_2
0.487 0.553 0.497 0.524
blood_val_3
0.532 0.574 0.512 0.530
blood_val_4
0.764 0.836 0.728 0.768
blood_val_5
0.519 0.615 0.538 0.614
blood_val_6
0.514 0.572 0.515 0.575
blood_val_7
0.510 0.578 0.529 0.534
blood_val_8
0.526 0.579 0.519 0.535
Main
Random
ASW Score
1.2k
10k
B
blood_val_1
0.553 0.642 0.355 0.419
blood_val_2
0.487 0.553 0.291 0.317
blood_val_3
0.532 0.574 0.382 0.416
blood_val_4
0.764 0.836 0.369 0.415
blood_val_5
0.519 0.615 0.323 0.390
blood_val_6
0.514 0.572 0.309 0.349
blood_val_7
0.510 0.578 0.386 0.417
blood_val_8
0.526 0.579 0.411 0.407
Main
OnlyValueEmb
C
blood_val_1
0.553 0.642 0.525 0.661
blood_val_2
0.487 0.553 0.470 0.501
blood_val_3
0.532 0.574 0.508 0.551
blood_val_4
0.764 0.836 0.750 0.809
blood_val_5
0.519 0.615 0.529 0.644
blood_val_6
0.514 0.572 0.494 0.531
blood_val_7
0.510 0.578 0.501 0.551
blood_val_8
0.526 0.579 0.504 0.525
Main
BinCls
D
ASW Score
Main 0.553 0.487 0.532 0.764 0.519 0.514 0.510 0.526
Only GEPC 0.472 0.406 0.467 0.738 0.477 0.454 0.460 0.475
Only MVP 0.518 0.483 0.509 0.775 0.512 0.493 0.516 0.515
Only TOPO 0.281 0.294 0.362 0.339 0.291 0.281 0.391 0.349
w/o GEPC 0.473 0.476 0.464 0.740 0.493 0.471 0.496 0.509
w/o MVP 0.495 0.482 0.467 0.717 0.488 0.470 0.484 0.509
w/o TOPO 0.497 0.433 0.500 0.759 0.539 0.477 0.523 0.500
blood_val_1 blood_val_2 blood_val_3 blood_val_4 blood_val_5 blood_val_6 blood_val_7 blood_val_8
0.7
0.6
0.5
0.4
0.3

**Figure S9. Ablation analysis of RegFormer variants across blood validation datasets.**

**(A)** Comparison between GRN-guided (Main) and Random models.

**(B)** Comparison between Main and OnlyValueEmb variants.

**(C)** Comparison between Main and BinCls variants.

**(D)** Heatmap of ASW scores across datasets for models trained with or without specific pretraining objectives (MVP, TOPO, GEPC).

### Figure S10

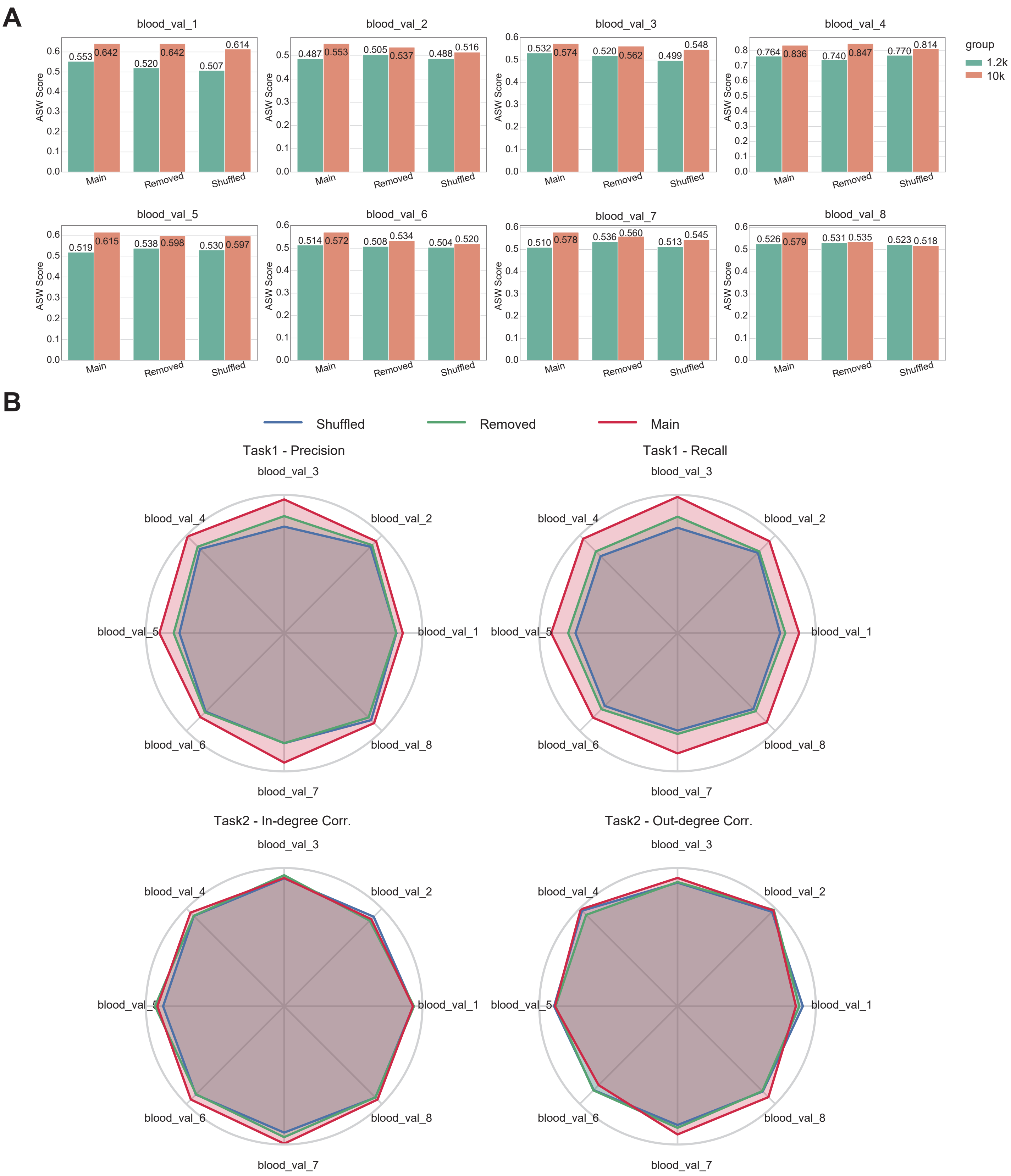


**Figure S10. Ablation analysis of GRN topology perturbations.**

**(A)** ASW scores across eight blood validation datasets comparing the Main, Removed, and Shuffled graph variants.

**(B)** Radar plots summarizing Task 1 (Precision, Recall) and Task 2 (In-degree and Out-degree Correlation) performance across datasets.

### Figure S11

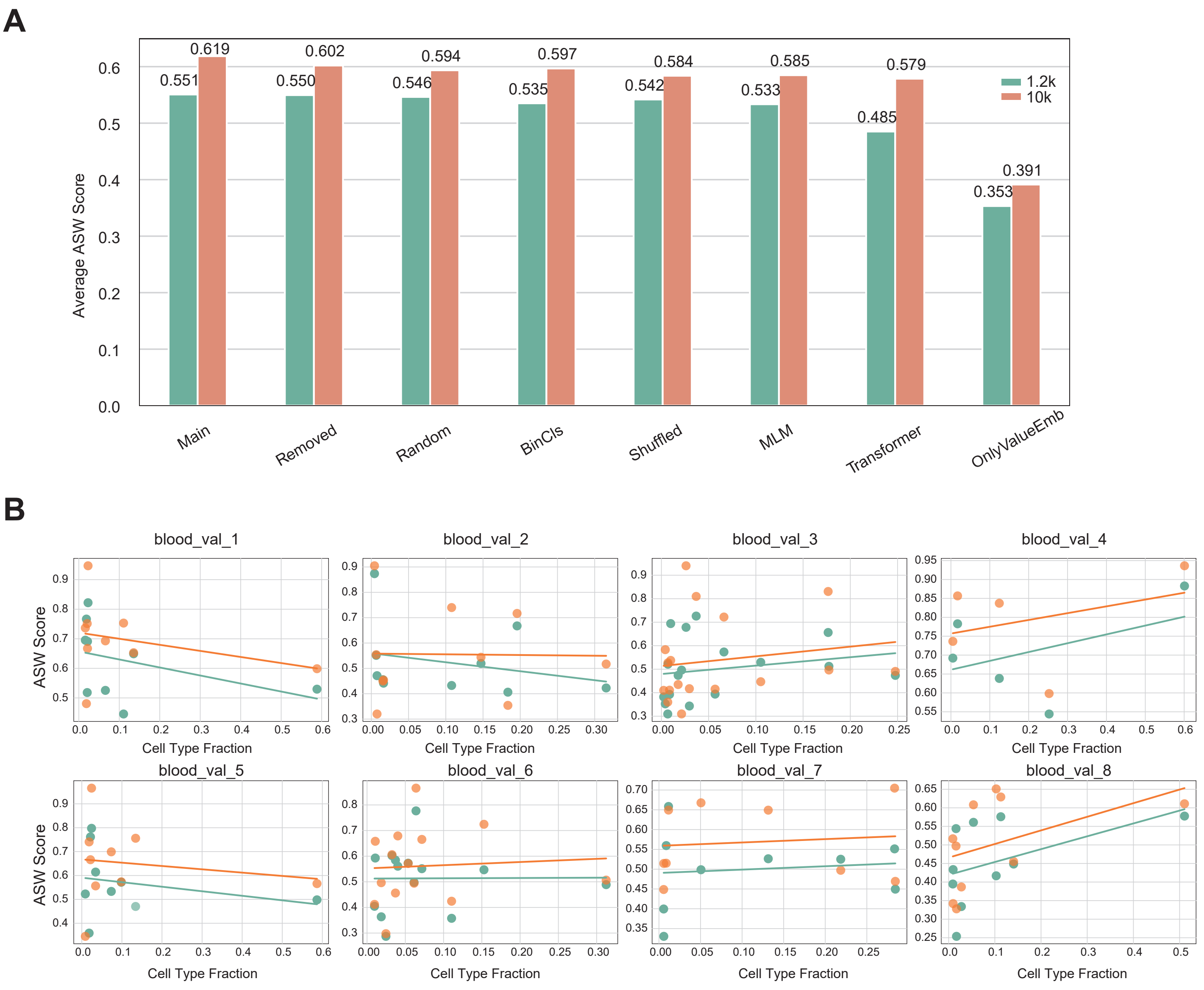


**Figure S11. Comparison of RegFormer performance at different sequence lengths.**

**(A)** Average ASW scores of RegFormer variants trained with sequence lengths of 1.2k and 10k.

**(B)** Relationship between ASW scores and cell type fractions across eight blood validation datasets for both sequence lengths.

#### Figure S12

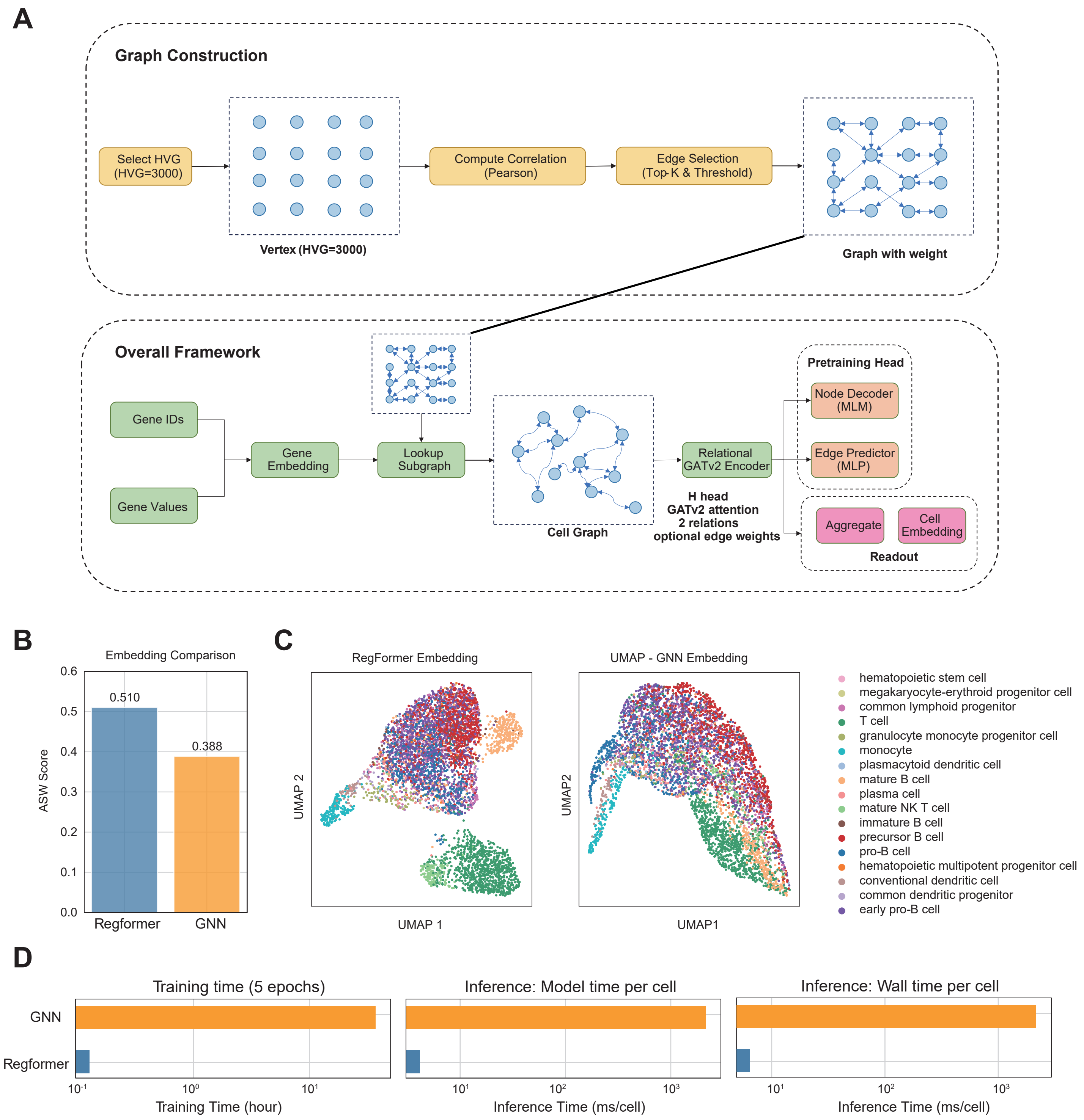


**Figure S12. Framework comparison between RegFormer and GNN for cell embedding.**

**(A)** Schematic of graph construction and overall modeling framework for both approaches.

**(B)** ASW score comparison between RegFormer and GNN embeddings.

**(C)** UMAP visualizations showing embedding structures from RegFormer and GNN.

**(D)** Training and inference time comparisons, including model and wall time per cell.

### Figure S13

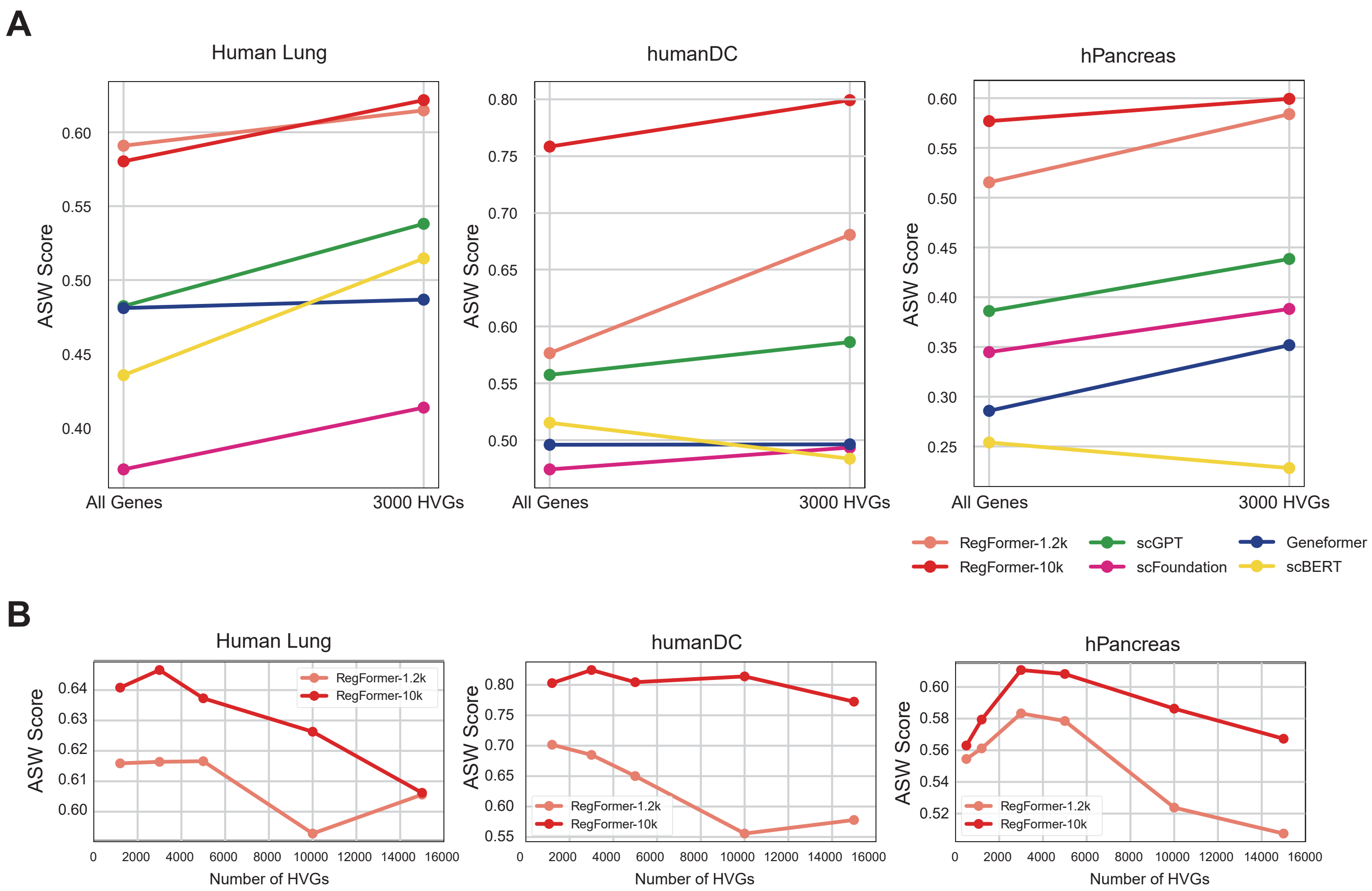


**Figure S13. Effect of highly variable gene (HVG) selection on model performance.**

**(A)** Comparison of ASW scores using all genes versus 3,000 HVGs across human lung, humanDC, and hPancreas datasets for different single-cell foundation models.

**(B)** ASW scores of RegFormer-1.2k and RegFormer-10k across varying numbers of HVGs.

### Figure S14

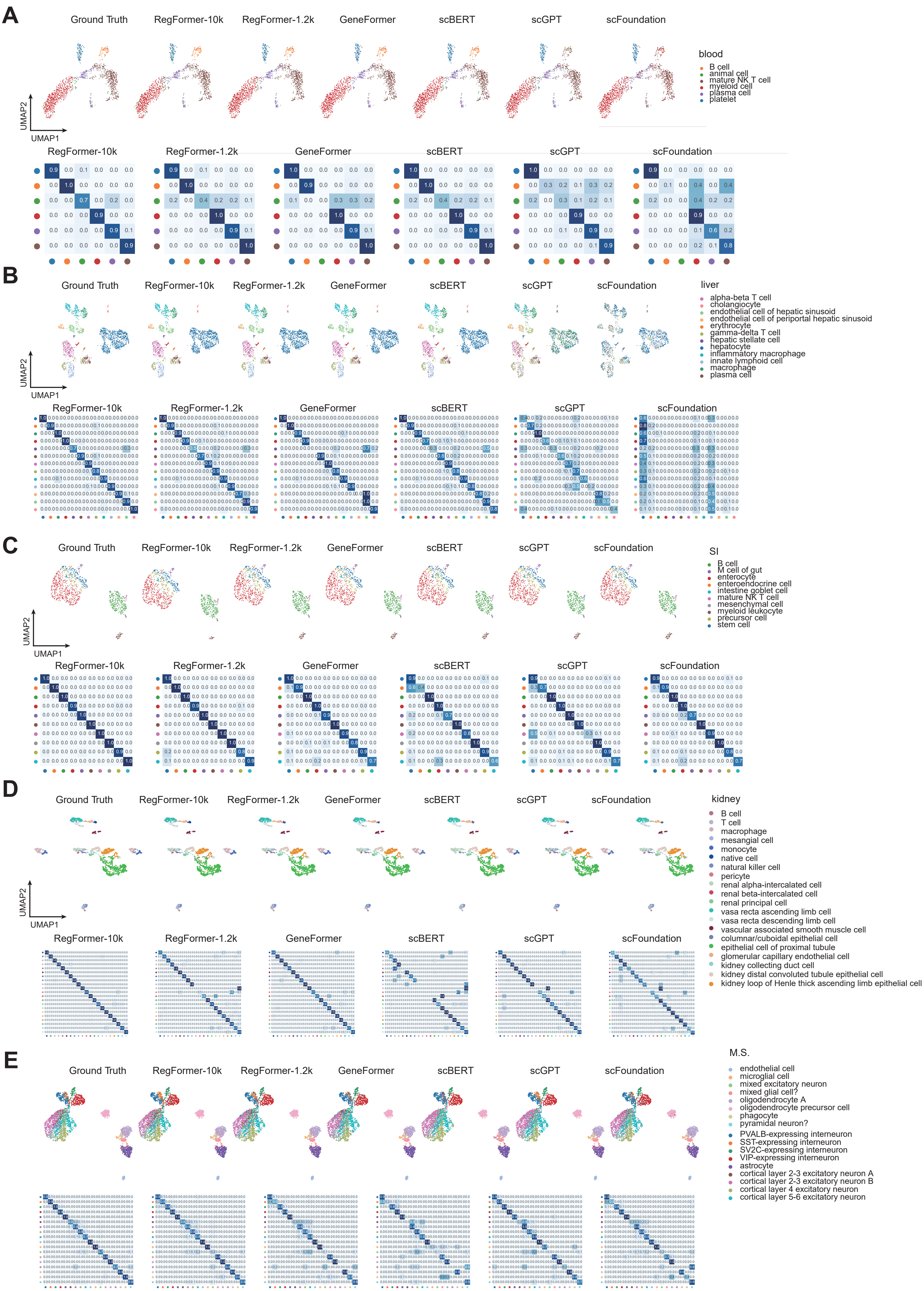
A
Ground Truth
RegFormer-10k
RegFormer-1.2k
GeneFormer
scBERT
scGPT
scFoundation
blood
B cell
animal cell
mature NK T cell
myeloid cell
plasma cell
platelet
UMAP2
UMAP1
B
liver
alpha-beta T cell
cholangiocyte
endothelial cell of hepatic sinusoid
endothelial cell of periportal hepatic sinusoid
erythrocyte
gamma-delta T cell
hepatic stellate cell
hepatocyte
inflammatory macrophage
innate lymphoid cell
macrophage
plasma cell
C
SI
B cell
M cell of gut
enterocyte
enteroendocrine cell
intestine goblet cell
mature NK T cell
mesenchymal cell
myeloid leukocyte
precursor cell
stem cell
D
kidney
B cell
T cell
macrophage
mesangial cell
monocyte
native cell
natural killer cell
pericyte
renal alpha-intercalated cell
renal beta-intercalated cell
renal principal cell
vasa recta ascending limb cell
vasa recta descending limb cell
vascular associated smooth muscle cell
columnar/cuboidal epithelial cell
epithelial cell of proximal tubule
glomerular capillary endothelial cell
kidney collecting duct cell
kidney distal convoluted tubule epithelial cell
kidney loop of Henle thick ascending limb epithelial cell
E
M.S.
endothelial cell
microglial cell
mixed excitatory neuron
mixed glial cell?
oligodendrocyte A
oligodendrocyte precursor cell
phagocyte
pyramidal neuron?
PVALB-expressing interneuron
SST-expressing interneuron
SV2C-expressing interneuron
VIP-expressing interneuron
astrocyte
cortical layer 2-3 excitatory neuron A
cortical layer 2-3 excitatory neuron B
cortical layer 4 excitatory neuron
cortical layer 5-6 excitatory neuron

**Figure S14. Multi-dataset cell type annotation comparison across models.**

**(A – E)** UMAP visualizations and confusion matrices showing predicted versus true cell types for blood, liver, small intestine (SI), kidney, and M.S. datasets, comparing RegFormer-10k, RegFormer-1.2k, Geneformer, scBERT, scGPT, and scFoundation.

#### Figure S15

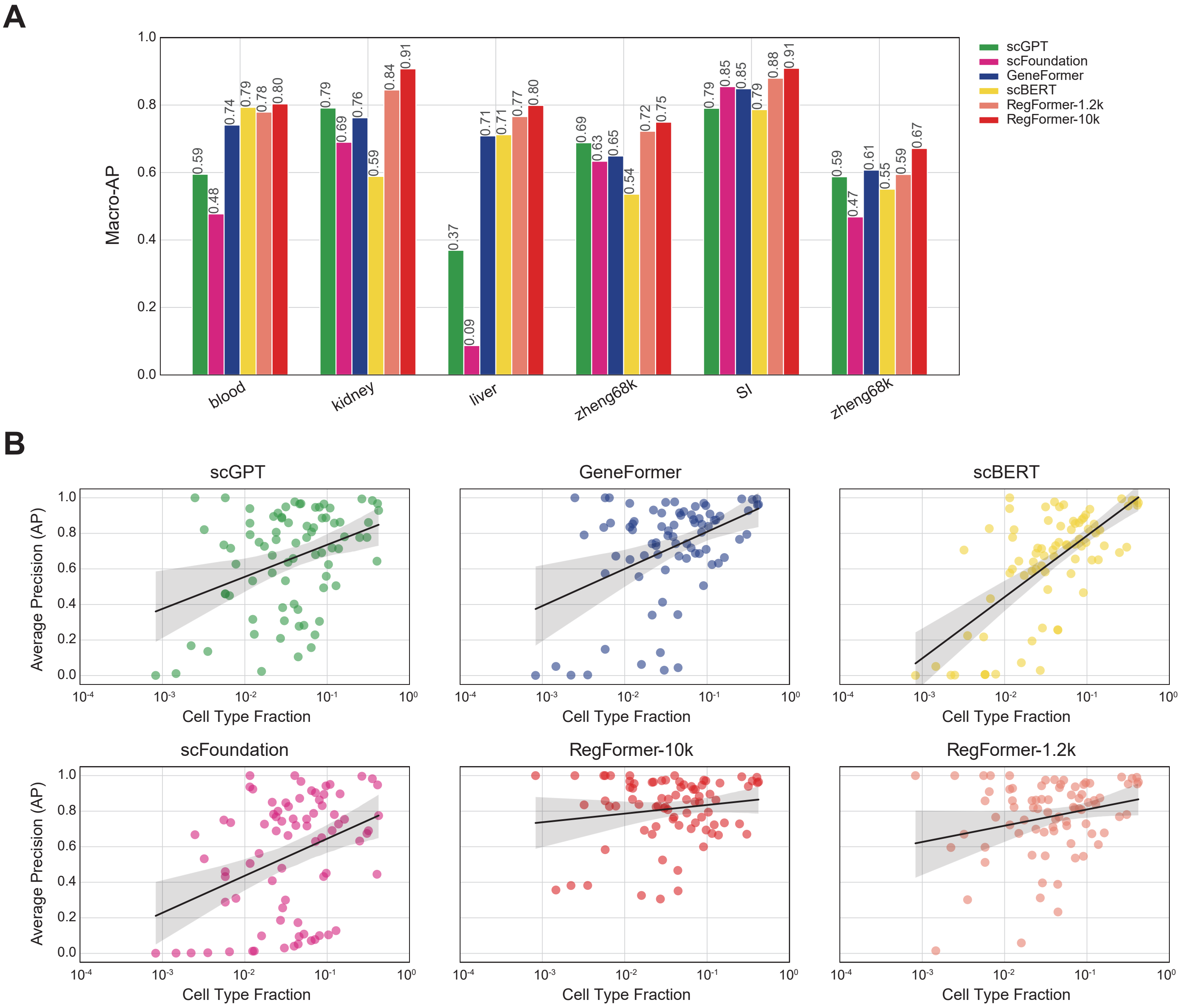


**Figure S15. Evaluation of macro-average precision (AP) across models and datasets.**

**(A)** Macro-AP scores of RegFormer and other single-cell foundation models across multiple datasets.

**(B)** Relationship between average precision and cell type fraction for each model.

### Figure S16

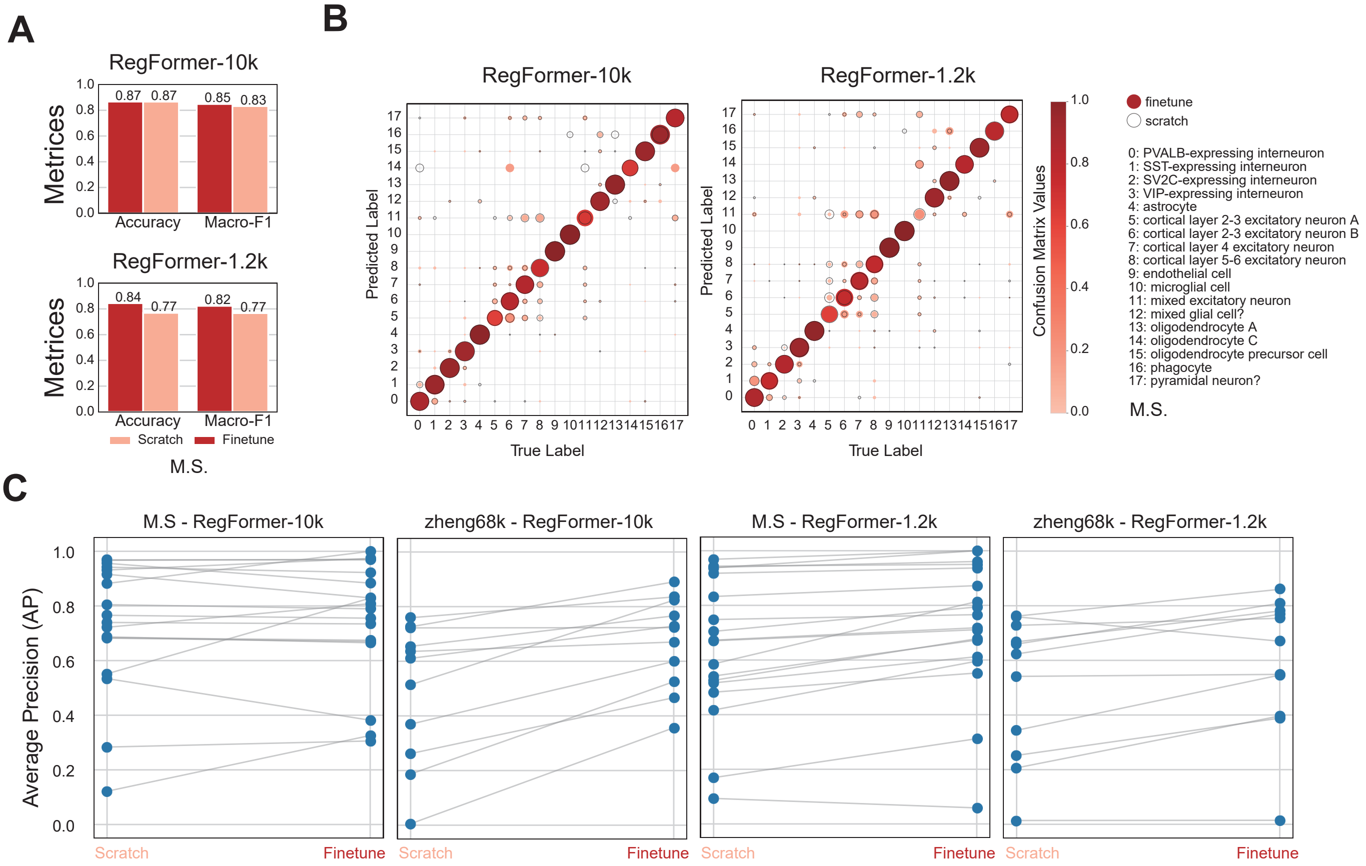


**Figure S16. Finetuning performance of RegFormer on cell type annotation tasks.**

**(A)** Accuracy and Macro-F1 comparison between finetuned and scratch-trained RegFormer-10k and RegFormer-1.2k models on the M.S. dataset.

**(B)** Confusion matrices showing predicted versus true labels for both training settings.

**(C)** Average precision (AP) comparison between finetuned and scratch models for each cell type across the M.S. and Zheng68k datasets.

### Figure S17

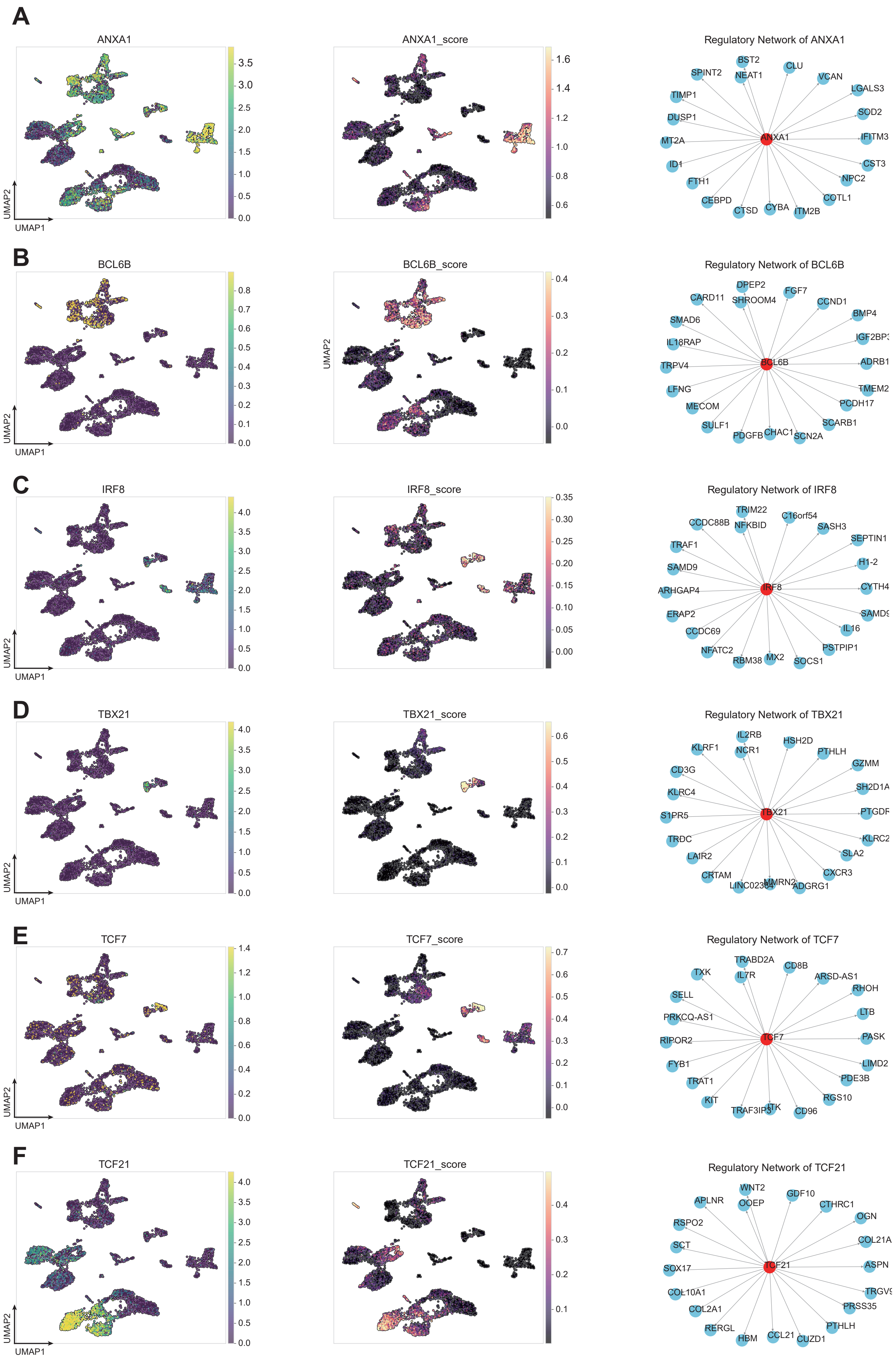

A
ANXA1
ANXA1_score
Regulatory Network of ANXA1
ANXA1
BST2
CLU
SPINT2
NEAT1
VCAN
LGALS3
TIMP1
SOD2
DUSP1
IFITM3
MT2A
CST3
ID1
NPC2
FTH1
COTL1
CEBPD
CTSD
CYBA
ITM2B
B
BCL6B
BCL6B_score
Regulatory Network of BCL6B
BCL6B
DPEP2
FGF7
CARD11
SHROOM4
CCND1
BMP4
SMAD6
IL18RAP
TRPV4
ADRB1
LFNG
TMEM2
MECOM
PCDH17
SULF1
SCARB1
PDGFB
CHAC1
SCN2A
C
IRF8
IRF8_score
Regulatory Network of IRF8
IRF8
TRIM22
C16orf54
CCDC88B
NFKBID
SASH3
SEPTIN1
TRAF1
H1-2
SAMD9
CYTH4
ARHGAP4
ERAP2
IL16
CCDC69
PSTPIP1
NFATC2
RBM38
MX2
SOCS1
D
TBX21
TBX21_score
Regulatory Network of TBX21
TBX21
IL2RB
HSH2D
KLRF1
NCR1
PTHLH
GZMM
CD3G
SH2D1A
KLRC4
S1PR5
TRDC
KLRC2
LAIR2
SLA2
CRTAM
CXCR3
MMRN2
ADGRG1
E
TCF7
TCF7_score
Regulatory Network of TCF7
TCF7
TRABD2A
CD8B
TXK
IL7R
ARSD-AS1
RHOH
SELL
LTB
PRKCQ-AS1
RIPOR2
PASK
FYB1
LIMD2
TRAT1
PDE3B
KIT
RGS10
ITK
CD96
F
TCF21
TCF21_score
Regulatory Network of TCF21
TCF21
WNT2
GDF10
APLNR
OOEP
CTHRC1
OGN
RSPO2
SCT
SOX17
ASPN
COL10A1
COL2A1
PRSS35
RERGL
PTHLH
HBM
CCL21
CUZD1
UMAP1
UMAP2

**Figure S17. Visualization of transcription factor (TF) regulons on the human lung datasets.**
**(A-F)** UMAP plots showing TF expression (left) and regulon activity scores (middle), along with inferred TF–target regulatory subnetworks (right) for representative TFs including ANXA1, BCL6B, IRF8, TBX21, TCF7, and TCF21.

### Figure S18

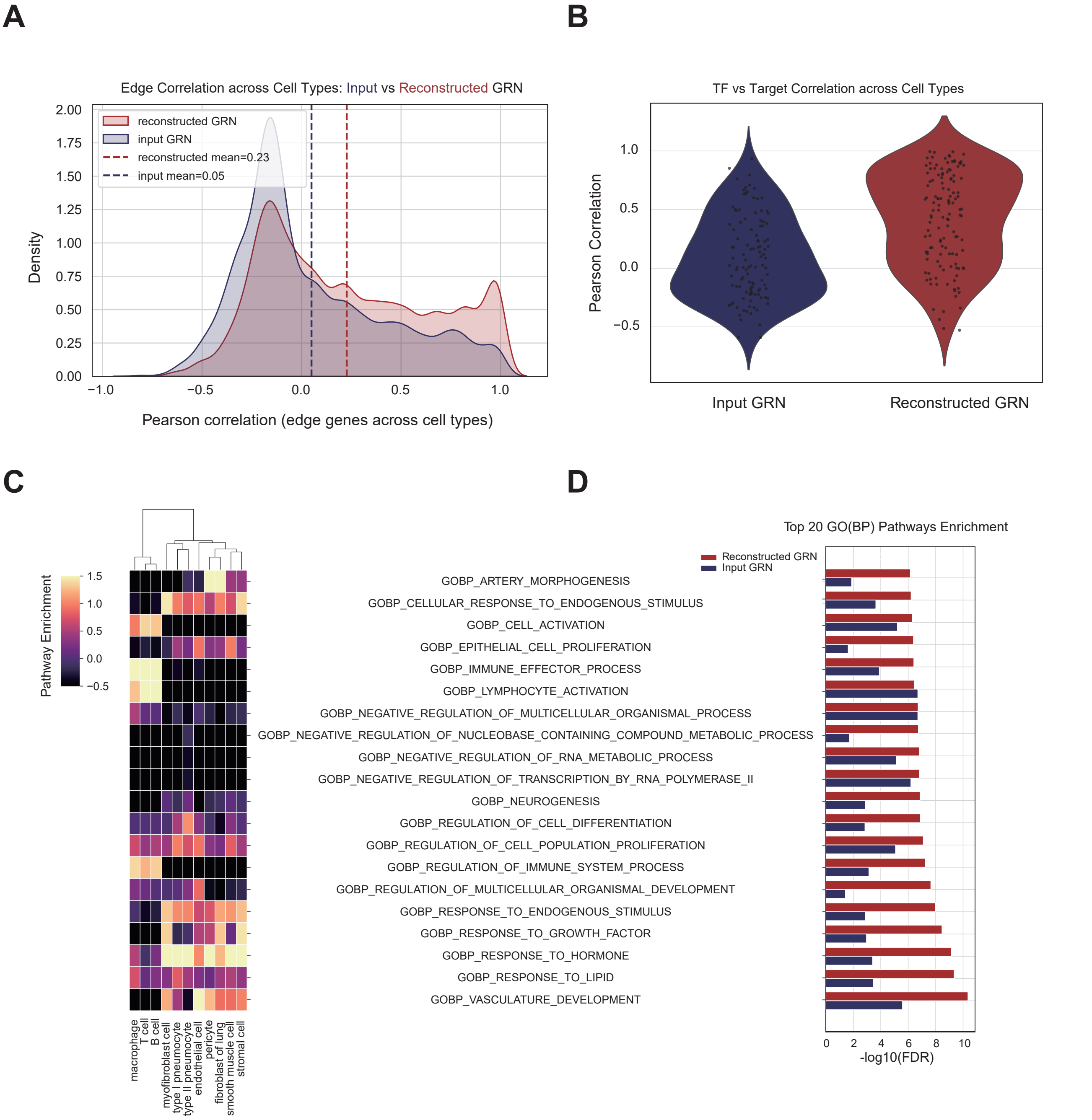


**Figure S18. Comparison between input and reconstructed gene regulatory network (GRN).**

**(A)** Density plots of edge correlations across cell types for input versus reconstructed GRN.

**(B)** Pearson correlations between transcription factors (TFs) and target genes across cell types for both networks.

**(C)** Heatmap of pathway enrichment across cell types in input and reconstructed GRN.

**(D)** Top 20 enriched GO Biological Process (BP) pathways comparing input and reconstructed GRN.

### Figure S19

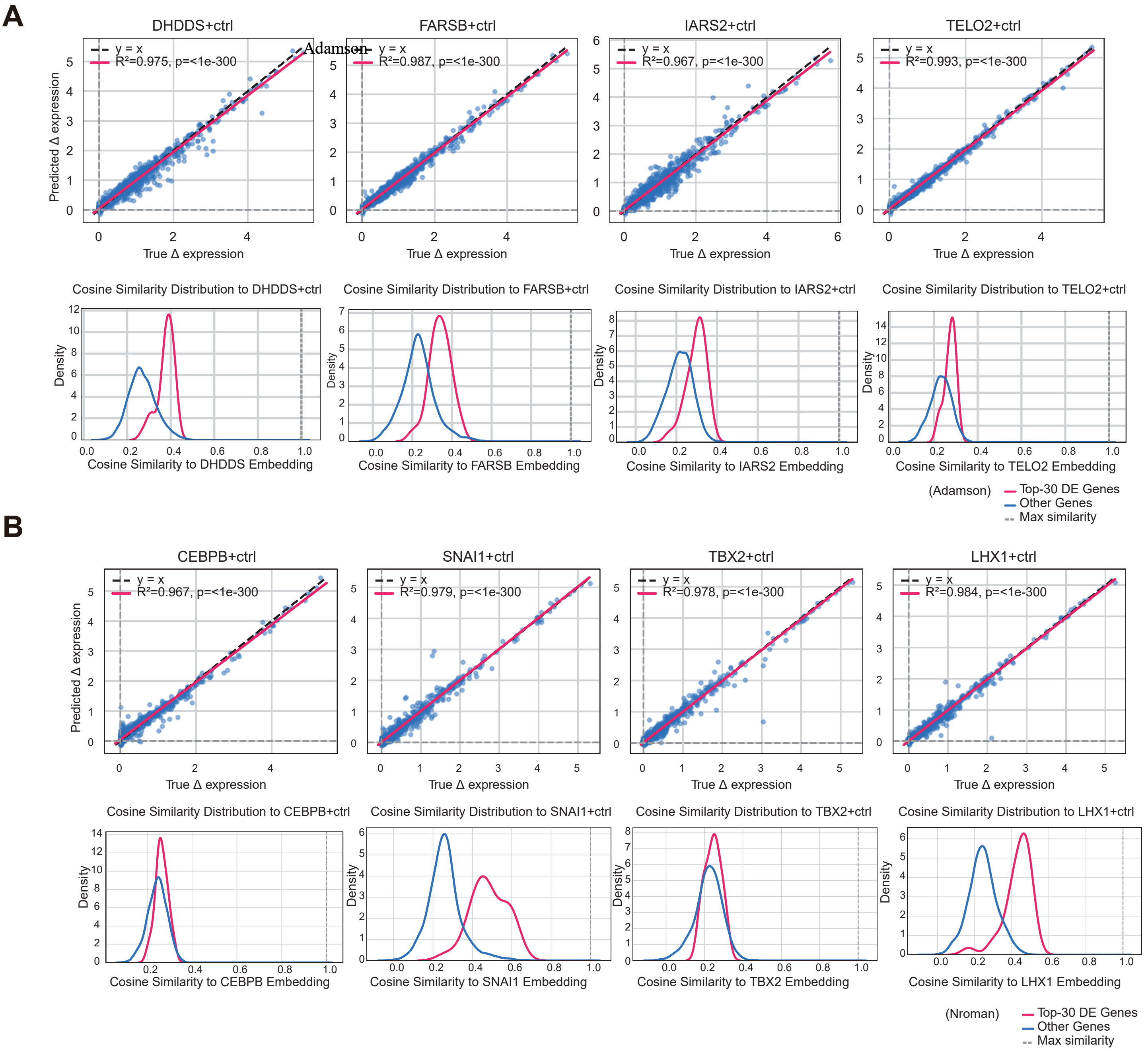


**Figure S19. Evaluation of perturbation prediction on the Adamson and Norman datasets.**

**(A-B)** Scatter plots of predicted versus true Δ expression and cosine similarity distributions between transcription factors (TFs) and differentially expressed (DE) genes for representative perturbations from the Adamson dataset (DHDDS, FARSB, IARS2, TELO2) and the Norman dataset (CEBPB, SNAI1, TBX2, LHX1).

### Figure S20

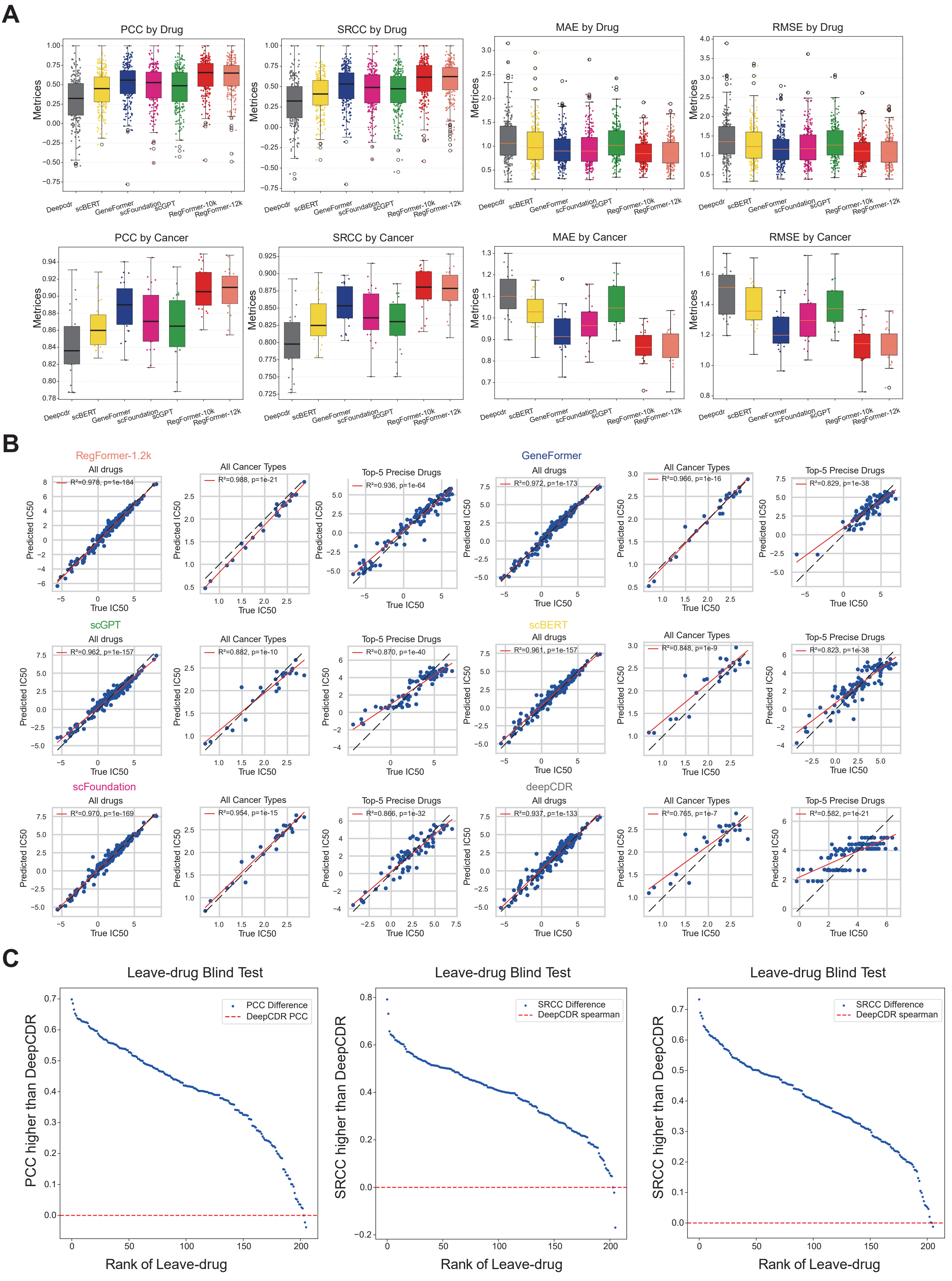


**Figure S20. Drug response prediction performance across models and evaluation settings.**

**(A)** Comparison of PCC, SRCC, MAE, and RMSE across drugs and cancer types for RegFormer and other single-cell foundation models.

**(B)** Scatter plots of predicted versus true IC50 values across all drugs, cancer types, and top-5 precise drugs for each model.

**(C)** Leave-drug blind test showing PCC and SRCC improvements of RegFormer over DeepCDR across ranked drugs.
